# Touching the (almost) untouchable: a minimally-invasive workflow for microbiological and biomolecular analyses of cultural heritage objects

**DOI:** 10.1101/2023.04.11.536414

**Authors:** Cecilia G. Flocco, Anika Methner, Franziska Burkart, Alicia Geppert, Jörg Overmann

## Abstract

Microbiological and biomolecular approaches to cultural heritage research have expanded the established research horizon -from the prevalent focus on the cultural object’s conservation and human health protection to the relatively recent applications to provenance inquiry and assessment of environmental impacts on heritage objects in a global context of a changing climate. Standard microbiology and molecular biology methods were developed for other materials, specimens, disciplines and contexts. Although they could in principle be applied to cultural heritage research, certain characteristics common to several heritage objects – such as uniqueness, fragility, high value and restricted access, demand tailored approaches. In addition, samples from heritage objects often yield low microbial biomass, rendering them highly susceptible to cross-contamination. Therefore, dedicated methodology that addresses these material limitations and operational hurdles along all procedural steps are needed. Here were propose a step-by-step standardized laboratory and bioinformatic workflow to analyse the microbiome of cultural heritage objects. The methodology was developed targeting the challenging side of the spectrum of cultural heritage objects, such as the delicate written record, while retaining modularity and flexibility to adapt and/or upscale the proposed workflow to heritage artefacts of a more robust nature or larger dimensions. We hope this review and workflow will facilitate the interdisciplinary inquiry and interactions among the cultural heritage research community.

## 1 Introduction

Over the last decades, the interest in the microbiome and the biomolecular make up of cultural heritage objects has significantly increased. The questions asked range from the longstanding focus on heritage conservation efforts and human health protection (Zielinska-Jankiewicz et al., 2008; Guiamet et al., 2011; Pinheiro et al. 2019) to the relatively recent focus on provenance inquiry and the study of the environmental factors impacting cultural heritage and related infrastructure in a context of a changing climate (Komatsu et al., 2019; West et al., 2019; Glevittzky et al., 2021; Pyzik et al., 2021)

For other objects of study or disciplines a section, portion or aliquot of the specimen or material of interest can be retrieved and used for microbiological and biomolecular analyses, such as blood or sputum for clinical investigation (Haldar et al., 2020; D’Aquila et al., 2021), soil, water or plant material for environmental studies (Lee et al, 2016; Vieira et al., 2020) or a core of a bone or a dental piece from human remains for archaeological and forensic research (Rascovan et al., 2016; Emmons et al., 2020). However, this is rarely possible in cultural heritage research, since the integrity of the object is of paramount importance. Certain characteristics inherent to most cultural heritage objects, such as their uniqueness, fragility, restricted or banned access (to the object or its location) and high value, among other factors, considerably hinder or preclude the possibility of gaining an actual portion of material for analysis (Prieto-Taboada et al., 2014). Therefore, dedicated, minimally-invasive sampling methods for cultural heritage research are required and need to be developed.

Although the choice of the sampling methodology will primarily be shaped by the characteristics of the object and the research questions to be answered, a surface sample is often the only viable alternative (Abel, 2011; Quye and Strlič, 2019). The sample itself will be a thin layer of material that is picked up by the sampling device. As a consequence, the retrieved samples may carry low microbial biomass, rendering them highly susceptible to contamination and other confounding effects introduced during the sampling event and subsequent laboratory processing (Eisenhofer et al., 2019). One exception to this prevailing sampling situation can occur when fragments of the object under study can no longer be relocated to its original position within the object, due to an advanced deterioration status; then, those delocalized fragments can be ‘sacrificed’ to provide an actual piece of the object for analytical procedures (La Russa et al., 2009). In addition, cultural heritage samples may contain a mixture of materials, originating from different sources and with different aging degrees. These properties can interfere with downstream molecular biology analyses, for example, by inhibition of certain enzymatic steps or oxidation of target molecules and with microbiological cultivation procedures due to the presence of biocides used for conservation (Schieweck et al., 2007; Mull et al., 2015).

Therefore, an interdisciplinary approach which considers the history and materiality of the heritage objects and employs dedicated sampling and analytical procedures for addressing and mitigating the described challenges are needed (Flocco, 2021; Curran and Zimmermann, 2022). Limitations should be carefully considered and embedded into the early stages of the experimental design and tracked downstream, along the data analyses and interpretation stages (Quye and Strlič, 2019). In addition, the envisaged methodology should be modular, to enable adaptation to different research needs, and include several safe stop points. The sampling method itself should be portable and robust enough as to be implemented beyond a laboratory context, since usually an on-site sampling of cultural heritage objects (for example, at archives, museums, or outdoor locations in the case of monuments) must be carried out. The implementation of the sampling method should be straightforward and accessible to operators not trained in laboratory procedures, considering scenarios in which the objects can be accessed and manipulated only by their curators or custodians.

In this work, we identify the main challenges and pitfalls bound to microbiological and biomolecular analysis of cultural heritage objects. We propose mitigation strategies and present a detailed and modular workflow, with suitable options to cover different research goals as well as resources and infrastructure availability scenarios. Aiming to translate theory into implementation, we propose a step-by-step laboratory protocol to analyse the microbiome of cultural heritage objects, streamlined through the combination of literature research and own experimental work, and provide guidelines for bioinformatic analyses. The methodology was developed aiming at the challenging end of the spectrum of cultural heritage objects, the century-old written record. Such experimental strategy enables a safe operational space and flexibility to adapt or upscale the methodology to objects of a more robust nature or larger dimensions.

Although we do not aim to exhaustively cover the extensive literature, we hope this tailored review and proposed experimental workflow will serve as a guidance to reliably explore the microbiome of cultural heritage research and facilitate interdisciplinary exchanges among the cultural heritage research community.

## 2 Experimental Design in Cultural Heritage Research: an Interdisciplinary Dialogue

Cultural heritage objects are often studied with an interdisciplinary approach covering history, archaeology, forensics, chemistry, materials science, molecular biology and microbiology (Mazzocchi, 2019; Piñar and Sterflinger, 2021; Flocco, 2021). In particular, the investigation of the biological materials composing or bound to heritage objects, such as proteins and the microbiome, has identified novel biomolecular historical markers (Teasdale et al., 2015; Rosenbloom, 2021) and re-kindled questions on the philosophy and ethics realm (Inkpen, 2019; Laplane et al., 2019.; Pálsidottir et al., 2019; Lange et al., 2022).

Interdisciplinary approaches can result in the design of new types of experiments and yield novel and enriched perspectives. For example, in our research project MIKROBIB (2018) we have challenged the prevalent view of microorganisms as agents of disease and deterioration, and explored their biographical and biotechnological potential from a holistic philosophical, microbiological and cultural perspective. In addition to dialogue and information exchange, *on site* visits of the life sciences researchers to cultural heritage collections and conservation facilities, and *vice versa*, can provide insight into each disciplines’ lines of inquiry, research approaches, conservation practices and the handling of the heritage objects and microbiological collections, enabling improved experimental design and adapted logistical procedures (Tobi and Kampen, 2018). Based on the interdisciplinary goals and the type of information needed, the microbiological and biomolecular tools and analyses have to be selected and adjusted accordingly. The research on cultural heritage objects may aim toward hypothesis testing, which can be achieved through the establishment of case studies (for example, by comparing contrasting situations around objects of interest to challenge a given hypothesis), or be directed toward the discovery and building of a hypothesis, for which exploratory approaches are used.

Microbiome studies embrace both, cultivation-dependent studies, in which microbial cells are grown and isolated in the laboratory, and cultivation-independent approaches, in which the cell constituents like nucleic acids, proteins or metabolites are analysed. Cultivation-dependent studies comprise the sampling and subsequent cultivation of microorganisms in a set of cultivation media and conditions, eventually yielding purified microbial isolates for further in-depth analyses, such as physiological, morphological and genomic studies. The accessibility of pure cultures for laboratory assessment, for instance, allows to detect novel biochemical or metabolic features that cannot be deduced from genome sequences alone (Overmann et al., 2017). On the other hand, this method will necessarily be constrained to taxa able to grow under the provided laboratory cultivation conditions. The approach will hence exclude those microorganisms for which suitable growth conditions are not achieved (Kapinusova et al., 2023)

Cultivation-independent microbiome studies can overcome these limitations. In particular, whole nucleic acids extracted from a given sample provide insights into the composition of microbial communities, independently of the viability status of the cells and the localization of the genetic material (intra- and extracellular). It is possible to work with the whole genetical material extracted from the sample and bioinformatically reconstruct the genomes of the microbes contained in such a mixture, as it is done in the so called metagenomic approaches (Quince et al., 2017), or focus on certain informative regions of the microbial genomes to obtain fingerprints of the different species present. The latter approach comprises marker genes, for example those codifying for certain key proteins, or non-protein-coding genes which can provide information on the identity of the microbial cells. One of the most widely used taxonomic markers is the gene encoding the RNA component of the 30 S subunit of the prokaryotic ribosome, the 16S rRNA gene (Woese and Fox, 1977; Woese et al., 1990). To increase the signal of these marker genes in a complex sample, the section of interest is amplified in the laboratory by means of a polymerase chain reaction (PCR) and specific primers. The resulting products, called amplicons, are further sequenced and analysed with bioinformatic tools. The 16S rRNA gene possesses highly conserved regions, which are targeted for PCR primer design, and hypervariable regions, which are informative of the identity of microbial entities. The 16S rRNA amplicon strategy can provide a taxonomic resolution power up to genus or down to species level, in some cases (Lundberg et al., 2013). The 16S rRNA amplicon strategy coupled to the increased accessibility of high throughput sequencing technology, provides a broad microbial community overview and readily detects microorganisms that escape cultivation (Gilbert et al., 2014). Thus, a combination of both cultivation and cultivation-independent assessments can provide a broader picture of the microbiome of cultural heritage objects since complementary information can be gained. A schematic representation of an interdisciplinary framework to studying the microbiome of cultural heritage objects is presented in Figure 1.

**Figure 1:**
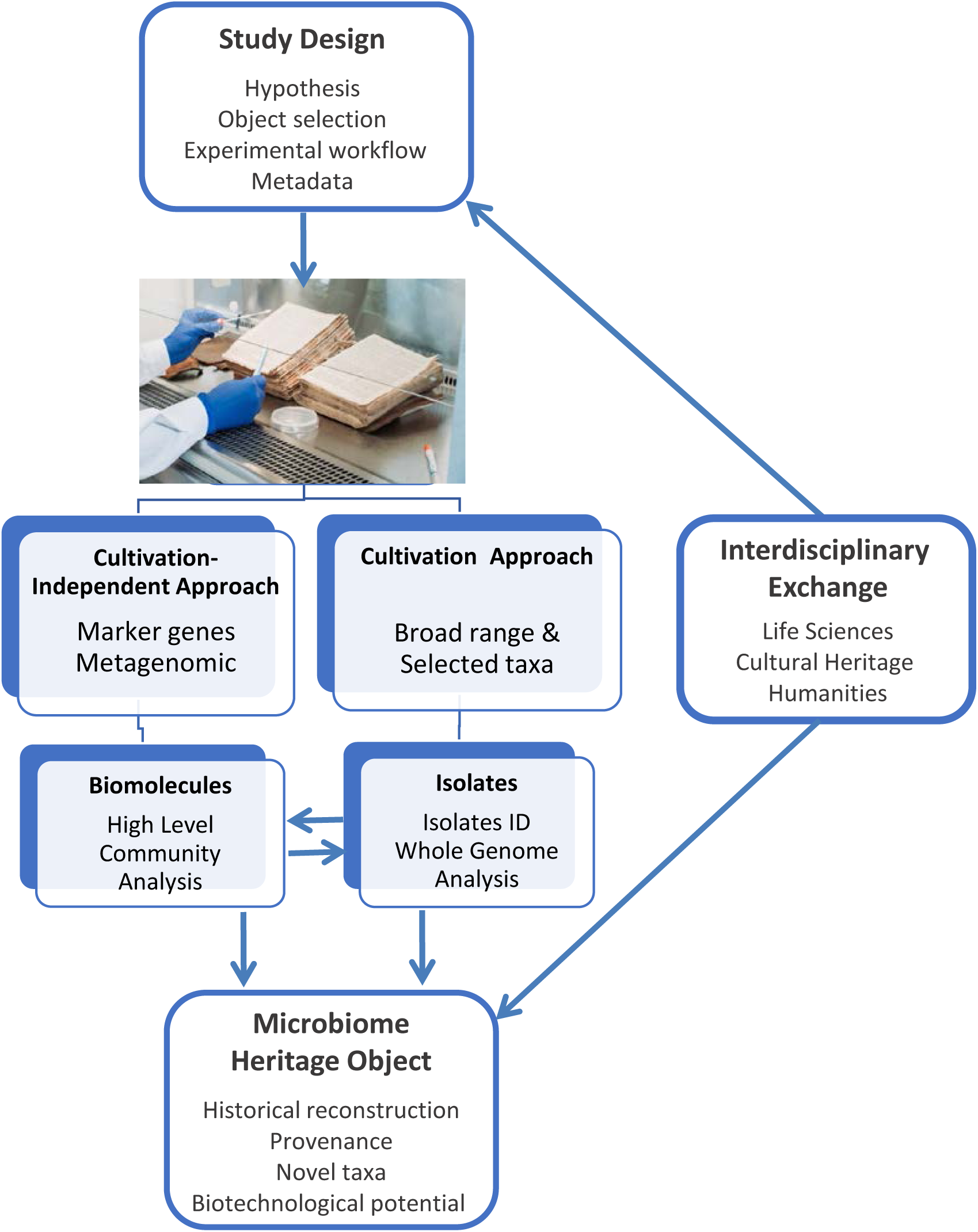
Schematic representation of an interdisciplinary framework to studying the microbiome of cultural heritage objects. Picture credit: C. Flocco, Leibniz Institute DSMZ.

Other possible analyses may target the materiality of the heritage objects. For example, the determination of the animal source for the parchment used for manuscripts is based on the analysis of the collagen proteins of the animal’s skin by mass spectrometry, coupled to a zoological database (Teasdale, et al., 2015; Brandt et al., 2018). Other examples are the determination of the type of pigments, binding agents and other materials used for the manufacture of heritage objects (Radini et al., 2019; Schuetz et al., 2019) or the origin of stains and spills (Fiddyment et al., 2020; Warinner et al., 2022). In the present work, we focus on the microbial community analysis through cultivation and cultivation-independent approaches.

## 3 General Considerations and Guidelines

### 3.1 Words matter: non-invasive versus minimally-invasive sampling

Although both expressions are often used in the literature and heritage conservation practice without strictly delimiting the boundaries for the ‘invasiveness level’ of a sampling method (Prieto-Taboada et al., 2014; Chaban et al., 2022), we think that a clear distinction is necessary when embracing a microbiological and biomolecular perspective. Several sampling methods could be regarded *a priori* as non-invasive since they apparently do no leave macroscopic traces or alter the integrity of the object. For example, surface sampling methods are often regarded as minimally-invasive (when compared to destructive methods extracting an actual piece of material), which is a crucial aspect from the point of view of object conservation (Multari et al., 2022). However, from the microbiological and biomolecular point of view, a section of an object that has been surface-sampled cannot be accessed again, since the sampling event itself will alter the microbial community and its microenvironment therefore, a subsequent sampling will likely not reflect the original condition. Considering that perspective, such a method is invasive. For large heritage objects this might not represent a limitation, since other sampling areas with the same or very similar characteristics might be available, but it may constitute a severe constraint for objects of small size or very heterogeneous shape and composition, for which replication of samples might not be feasible. In this case, a careful examination of the object and its accompanying metadata prior to the sampling event is necessary to guarantee a representative sampling and avoid on-the-fly decision-making. When available, digitized versions of the objects under study (for example a manuscript or codex) are of great help for the experimental design and preparation of sampling, since it allows to explore the document in advance, without additional manipulation that may damage or contaminate the samples. This is of particular advantage if multiple analyses are to be implemented as part of an interdisciplinary project.

### 3.2 Access to objects

In most scenarios, sampling of heritage objects for microbiological or other analyses must be conducted *on site*, given that such objects are part of collections (for example, at museums and libraries) and cannot be displaced due to safety or insurance reasons or, given their dimensions and other properties, cannot not be mobilized (for example, monuments or historical buildings). Therefore, the sampling methodology should be portable, robust and easy to implement *on site* (Brunetti et al., 2016). In some cases, only dedicated personnel (custodians, curators) may have access to the objects of interest, therefore, clear instructions and protocols for all sampling steps and appropriate personal protective equipment should be provided, in order to take a representative sample and avoid contamination during manipulation. If the samples are to be processed *ex situ*, adequate sample preservation and storage, as well as suitable transport conditions (e.g., with refrigeration or preservation solutions) are needed.

### 3.3 Visible heterogeneity

Depending on their intrinsic characteristics, surrounding environment and use, cultural heritage objects can be very heterogeneous regarding their shape, structure, material composition and aging status. Therefore, treating the objects as a landscape or habitat with different zones or microenvironments can help to chart the heterogeneity of an object and aid the interpretation of experimental results. For example, in the case of the written heritage, it is possible to define areas impacted by the ink and pigments, that is, the text and illuminated areas, versus areas without scripture, such as the margins. Heavily touched areas, such as the corners or sides of the page, where fingerprints and skin oils are left, can harbour a microbial community that differs from those found in areas with lower touch impact. Also, what is actually written on certain pages or sections also matters, since the texts can provide hints to the use and history of an object (Flocco, 2021). An interdisciplinary exchange with the cultural heritage professionals can reveal which pages are likely to be more touched, as is the case for some religious books, since certain passages are visited more often than others. Other areas of the book can be shaped by different levels of exposure to the environment (humidity, dust, light), which will have a strong influence on the microbial communities (Glevitzky et al., 2021). Beyond the pages or folia, other materials to analyse are the codex or book covers, the type and materials used for binding, pollen grains, the holes and depositions made by insects, and the insects themselves, dust, wax from candles used for reading and traces of human secretions (Edwards and Munschi, 2005; Dallongeville et al., 2016 and references therein). These are some examples of the biomolecular make up of an object, accumulating and dynamically changing along the history of an object.

### 3.4 Heterogeneity under the biomolecular lens

Based on the history, use and materiality of an object elucidated through interdisciplinary exchanges, particular physicochemical conditions can be anticipated, providing guidelines for the design of microbiological studies and sampling procedures. For example, specific features bound to an object would impair microbiological cultivation, for instance, the treatment with biocides for conservation purposes, the presence of heavy metals arising from inks or metallic parts such as book clasps. Other materials can interfere with cultivation-independent steps, such as polyphenols and other humus-like substances present in plant based-inks, since they can bind to the nucleic acids and interfere with PCR amplification processes (Schrader et al., 2012). Figure 2 shows the aspect of a swab used for sampling a parchment codex examined by scanning electron microscopy (SEM), displaying some potential microbial structures surrounded by particles of unidentified debris.

**Figure 2:**
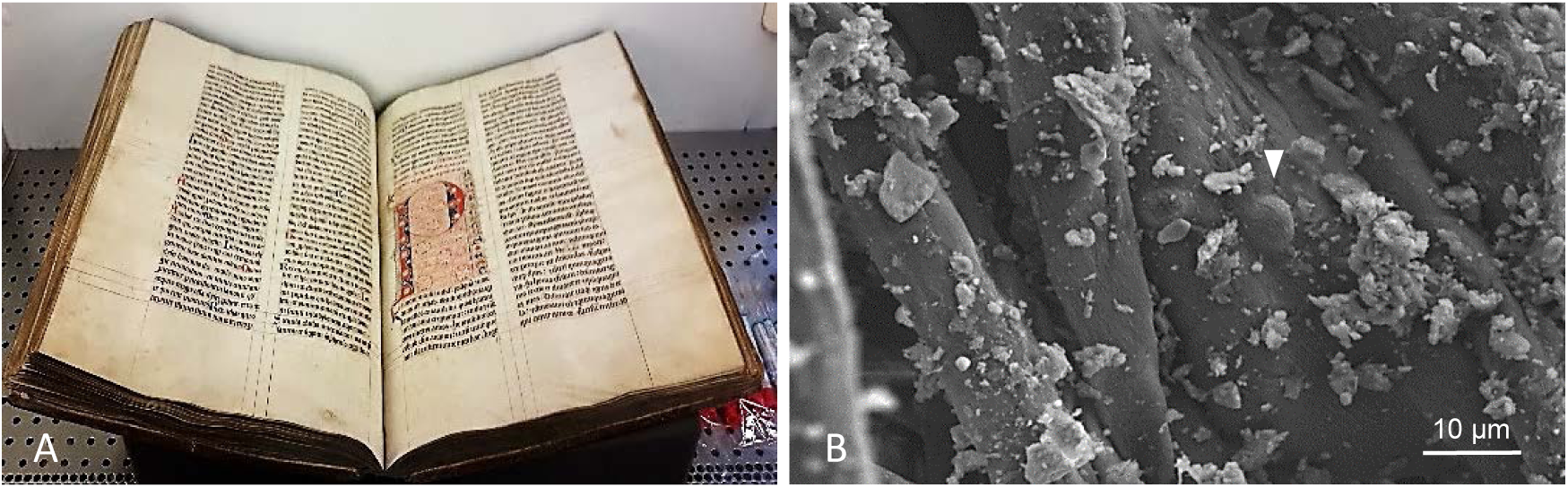
(A) Representative experimental set up for sampling a written cultural heritage object (Ms 12, a parchment codex) *on site* (at University Library Leipzig, UBL) under aseptic conditions, using the swabbing method. (B) Scanning electron micrograph showing a swab used for sampling the parchment codex (UBL, Ms 12). The arrow points to a possible endospore structure nested in the fibres of the swab head, surrounded by unidentified debris from the sampled object. Picture credits: (A) C. Flocco, Leibniz Institute DSMZ, in accordance with UBL. (B) C. Flocco, Leibniz Institute DSMZ and Manfred Rohde, Helmholtz Centre for Infection Research (HZI).

The microbial cells present in a cultural heritage object can be classified according to their source (intrinsic to the object, arising from users, deposited with dust particles, by insects, etc) and status (viable cells, viable but dormant, not viable). A similar scheme applies to the nucleic acids present in the sample (Teasdale et al. 2017) which, in addition can be categorized according to their location or cellular compartment (intracellular or extracellular) and age (contemporary to ancient, with different degrees of degradation; Vai et al., 2016)

Given the described macro- and micro-heterogeneity scenario, we suggest to take as many replicate samples as possible, to increase the chances of capturing most of the variability. Technical replicates are also recommended but priority should be given to experimental replicates when the material available for sampling is limited. In the latter case, it should be considered that certain samples may represent a unicate. Therefore, analytical procedures and results interpretation should be adapted accordingly, as we describe in the next sections.

### 3.5 Analytical challenges: low biomass, low diversity

Microbiome studies from cultural heritage objects are oftentimes particularly challenging in analytical terms, given a specific microbiological constellation: samples may carry low microbial biomass and/or low biological diversity. This is often the situation in which microbial colonization of a heritage object is not macroscopically visible. For example, very well conserved manuscripts or other cultural objects with dry and even surfaces offer relatively adverse conditions for microbial proliferation. Such potential scenario calls for a proper adjustment of the experimental design and all downstream steps of the workflow, from the sampling method to the sequencing and the microbial cultivation approaches. In addition, some *in silico* mitigation measures can be implemented *a posteriori*, by adjusting the parameters of the bioinformatic pipeline, but the adaptation of upstream processes should have priority.

The personnel involved in the sampling event itself and laboratory procedures should wear appropriate personal protection equipment to avoid the shedding of own biological material (for example, skin cells, saliva and the attached microbiota) on the objects and sampling materials. When possible, the sampling of mobile heritage objects should be carried out under a clean bench. Some libraries, archives and museums often have such in-house facility and it might be possible to bring in the object of study. When such facilities are not available, the sampling will be done *on site*, which increases the possibility of contaminating the samples. In this case, adequate mobile laboratory instruments should be considered, in addition to appropriate personal protection equipment, together with an *a priori* carefully designed sampling scheme to minimize unnecessary manipulations and exposure to spurious contamination.

Various protocols for the extraction of nucleic acids from diverse materials exist, but those usually require relatively large amounts of starting material and employ harsh extraction methods or lengthy processes, which can produce additional fragmentation of the DNA. The latter will, in turn, impact the sequencing step and the correct assembly of the generated amplification fragments, impairing downstream analyses. The sampling tool or method should maximize both, the *uptake* of the analyte of interest (microbial cells, nucleic acids) and the effective *release* of such material from the sampling device. Similarly, the methods for extraction of nucleic acids for cultivation-independent microbiome analyses should strike a balance between efficiency and product degradation protection, to minimize damage to the extracted nucleic acids (which may already present advanced fragmentation in the case of ancient materials). Typically, it should enable to break hard-to-lyse microbial cells, such as endospores, which is the status in which some bacterial species remain viable over time under unfavourable environmental conditions, while yielding high molecular weight products. Given that the nucleic acids extract may carry a mixture of different molecules that can hinder downstream processes, it is important to include purification steps, to reduce or eliminate such compounds. Eventually, such purification procedures can also be used to concentrate the nucleic acids extract by reducing the volume of the eluent reagent.

Low biomass samples have a very low signal to noise ratio, that is, the signal produced by the target analyte (total microbial nucleic acids, PCR amplicon signal or microbial counts, for example) can be very close to the signal spuriously produced by the instrument or measurement method or by contaminants introduced along the workflow (Kennedy et al., 2023). Therefore, it is very important to intensify the set of controls to enable the identification of microbial entities introduced into the sample as contamination. We recommend to include environmental controls (for example from the air surrounding the objects) as well as processes and reagents controls (not containing a sample from the object itself). This can be achieved by applying the same nucleic acids extraction method to the sampling devices (for example, swabs) that did not touch the object of interest, and then also run extraction reactions and other downstream procedures (such as PCR reactions and the sequencing library preparation) with the same reagents in the absence of sample. These controls enable the detection of contaminants in commercial kits and in-house prepared reagents or bound to the equipment (Salter et al., 2014). The sampling order is also of importance; we recommend to take environmental controls prior to taking the object’s samples to avoid the potential generation of clouds of particulate material that could produce a confounding effect between samples and controls. Additionally, we recommend using plastic labware that has reduced capacity to bind nucleic acids for all steps involving work with nucleic acids and derived PCR products. Bulk liquid reagents such as buffers and glassware can be treated under UV light to reduce possible contamination.

Further downstream in the experimental workflow, dedicated protocols for sequencing low biomass and/or low diversity samples should be implemented. In some cases, reagents and technology suppliers provide such dedicated protocols, in others those have to be developed and validated in house. Since the yield of the sequencing libraries can be very low, we emphasize the importance of using labware with low affinity for nucleic acids in order to minimize losses, and include purification and concentration steps, as previously mentioned. In addition to low microbial biomass, samples from cultural heritage objects may carry low microbial diversity. Such situation poses technical challenges for high throughput amplicon sequencing approaches, as it has been shown in studies dealing with low diversity samples of environmental and medical origin (Wu et al., 2015, Kennedy et al., 2023 and references therein). Low microbial diversity samples generate rather homogeneous 16S rRNA amplicon pools, a feature that hinders the initial sequencing steps requiring nucleotide diversity for efficient cluster identification and calibration procedures (Illumina, 2014; Shirmer et al., 2015). A strategy commonly used to solve this technical challenge is the addition of a high diversity sequencing library (typically a phage PhiX library) which is co-sequenced with the pool under study (Fadrosh et al., 2014). This modification increases the sequence diversity of the library pool and facilitates the correct execution of the mentioned initial sequencing steps. Also, the barcodes and indexes used to generate the sequencing construct add a degree of sequence diversity, contributing towards overcoming the technical issue (Kozich et al., 2013). The s manufacturers of sequencing platforms provide dedicated protocols and reagents to tackle low diversity samples (Illumina, 2014) but, as mentioned, in house optimization and tailored modifications might still be necessary.

## 4 Adjusting the Workflow: Pilot Studies

Since the experimental workflow for studying the microbiome of cultural heritage objects involves a large number of steps, it can be difficult to spot bottlenecks a *priori* or disentangle the source of a problem at the end of a long series of concatenated procedures. Therefore, it is recommended to run pilot studies, preferentially using objects of similar characteristics to the one under study (when available), or a section of the same object that can be assigned to this purpose. This approach allows identifying beforehand the potential limitations of each step of the protocol and adjust accordingly at that position of the workflow. Running such a pilot study might appear as an unnecessary additional burden at first sight. However, given the several pitfalls that can be encountered along the road and the uniqueness and high value of heritage objects, the approach can save both, time and resources in the long run.

In this section, we describe the main findings of our pilot study, in which we tested crucial steps of the experimental workflow. We used the gained information to design and optimize the step by step protocol provided together with this work. For purposes of comparability, we conducted all experiments on a non-catalogued parchment codex of the XIV century held at the Manuscript Center of Leipzig University Library (Ms 12; Leipzig, Germany), which was kindly made available for this purpose. Since the book could not be mobilized from its custody institution, the samples and initial microbiological processing were done at the conservation department of the Leipzig University Library (Figure 2A). The requisite for *on site* work provided the opportunity to test as well the portability and adaptability of the sampling procedures. The samples were properly prepared and transported to our labs (in Braunschweig, Germany) for conducting the subsequent microbiological and biomolecular research.

### 4.1 Choosing the sampling method

Selecting a sampling method for cultural heritage objects should be done considering the ethical sampling guidelines of the Institute of Conservation (ICON; Quye and Strlič, 2019), recommending to initiate the research using the least invasive, most accessible and cost-efficient sampling method.

Among all possible minimally invasive sampling procedures for written cultural heritage objects previously outlined, we focused on those that could be simultaneously used for both, cultivation and cultivation independent microbiome research. We shortlisted two surface sampling methods: membrane blotting and swabbing. Membrane blotting involves the use of a membrane (for example, cellulose nitrate membranes) which is placed on top of the surface for a determined period of time to collect the microbiota and subsequently analysed by microscopy and biomolecular approaches. This surface sampling method is suitable to materials with a very delicate surface coat, such as paintings, photographs or albumen prints (Puškárová et al., 2016). However, it can be difficult to adapt this method to objects that do not have a flat surface. Swabs are more versatile for surface sampling diverse materials and objects with different shapes. It should be considered that the materials composing both, the swab tip and stem, can impact downstream process (Wise et al., 2021).

We tested the membrane blotting method over swabs on a parchment codex by assessing the count of microbial colonies recovered after incubation in a range of solid cultivation media. The array on cultivation media was designed based on assumptions of the possible microorganisms present in a given sample, considering both the materiality of the object and descriptions of microbiological findings in libraries and archival material found in the literature (prevailingly bacteria and fungi). Although the microbial counts recovered from a single sampling situation were in a similar range for both methods (range: 0-15 colonies per plate per cultivation medium; Flocco and Overmann, 2021) we found that the membrane method may hinder the microbial cultivation survey, depending on the type of organisms present in the sample and their growth pattern. For example, we observed that biofilm-producing microorganisms produced a continuous growth line along the rim of the blotting membrane when placed on a solid cultivation medium.

This exuberant growth formed a continuous ring around the membrane and engulfed other microbial colonies in the area, as shown on Figure 3. This growth pattern impaired both, colony counting and the isolation of single microbial entities, therefore we discarded this method for this type of heritage object.

**Figure 3:**
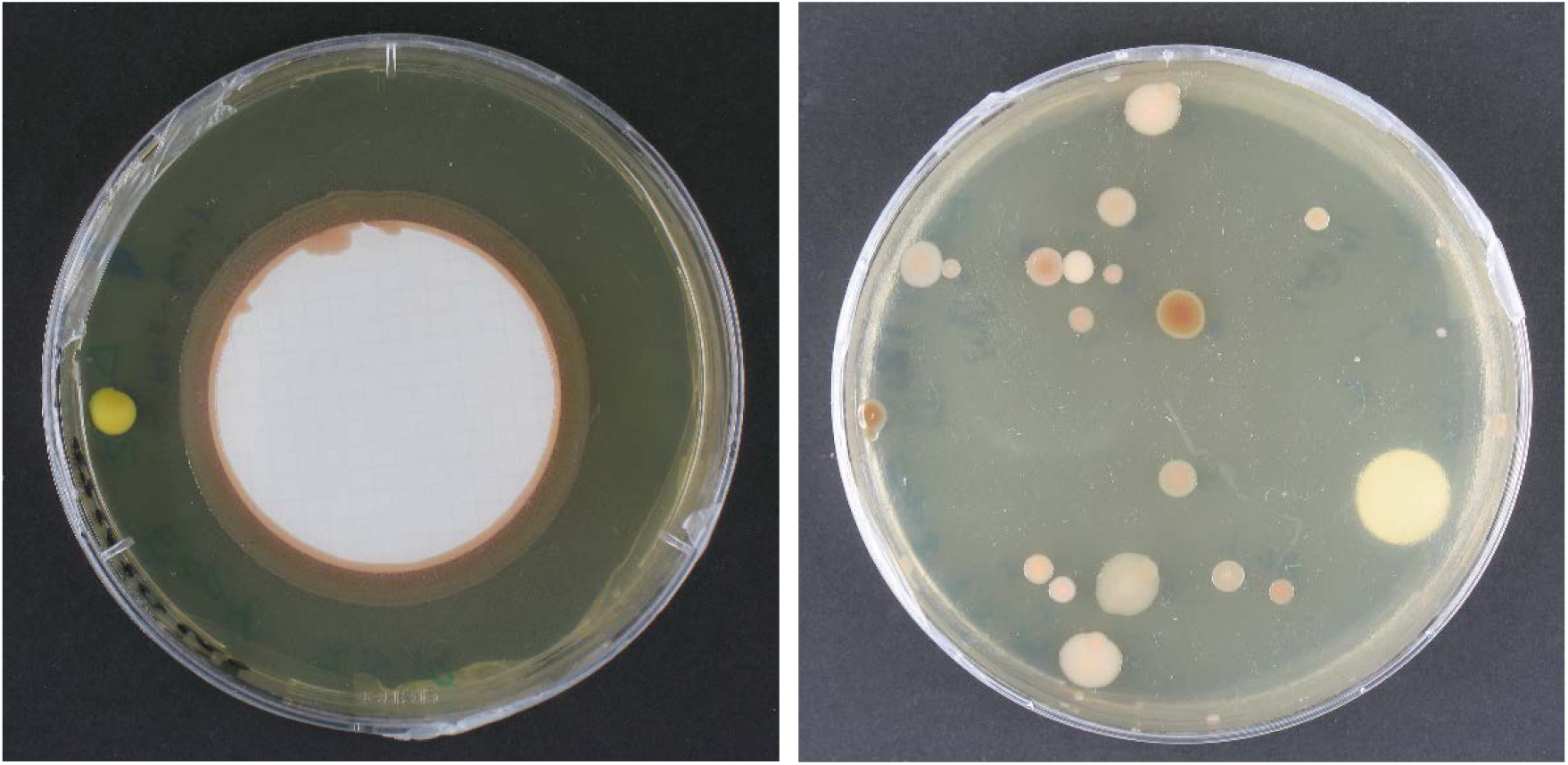
Comparison of the performance of the sampling methods on solid cultivation medium. A) Agar plate with the nitrocellulose membrane used for sampling, showing continuous growth along the micro-dent in the solid cultivation medium formed around the membrane border. B) Agar plate showing well defined and spaced microbial colonies obtained by direct spread plating with the swab used for sampling. Both samples were taken from comparative areas of a parchment codex (UBL, Ms12) and cultivated on trypticase soy yeast medium at room temperature (∼ 22 °C). Picture credit: C. Flocco, Leibniz Institute DSMZ.

As for the swab method, it is desirable that the device collects as much material as possible but then easily releases it during subsequent processing. We tested a series of standard cotton tip swabs and nylon flocked ones from different brands and measured the amount of nucleic acids that could be extracted from comparable surface samples of the parchment codex mentioned above. Our pilot assay showed that the nylon flocked swabs had a superior performance, when compared to swabs with other tip material (cotton and viscose; Figure 4). In addition, it should be considered that the cotton swabs may leave fibres on the objects, which could serve as a nutritional source for certain microbes, thus adding a potential risk to the manuscript. A second criterium for the selection of the sampling method emerged from the hands-on experience: the flocked swabs used in our study (Copan, Brescia, Italy) have a flexible stem and come with variable stem lengths, head size and geometry. These options allow adapting the same sampling tool to the different shapes, textures and intricate structures of cultural heritage objects while conserving the same sampling principle and performance since the swab tip design and functionality remains unchanged.

**Figure 4:**
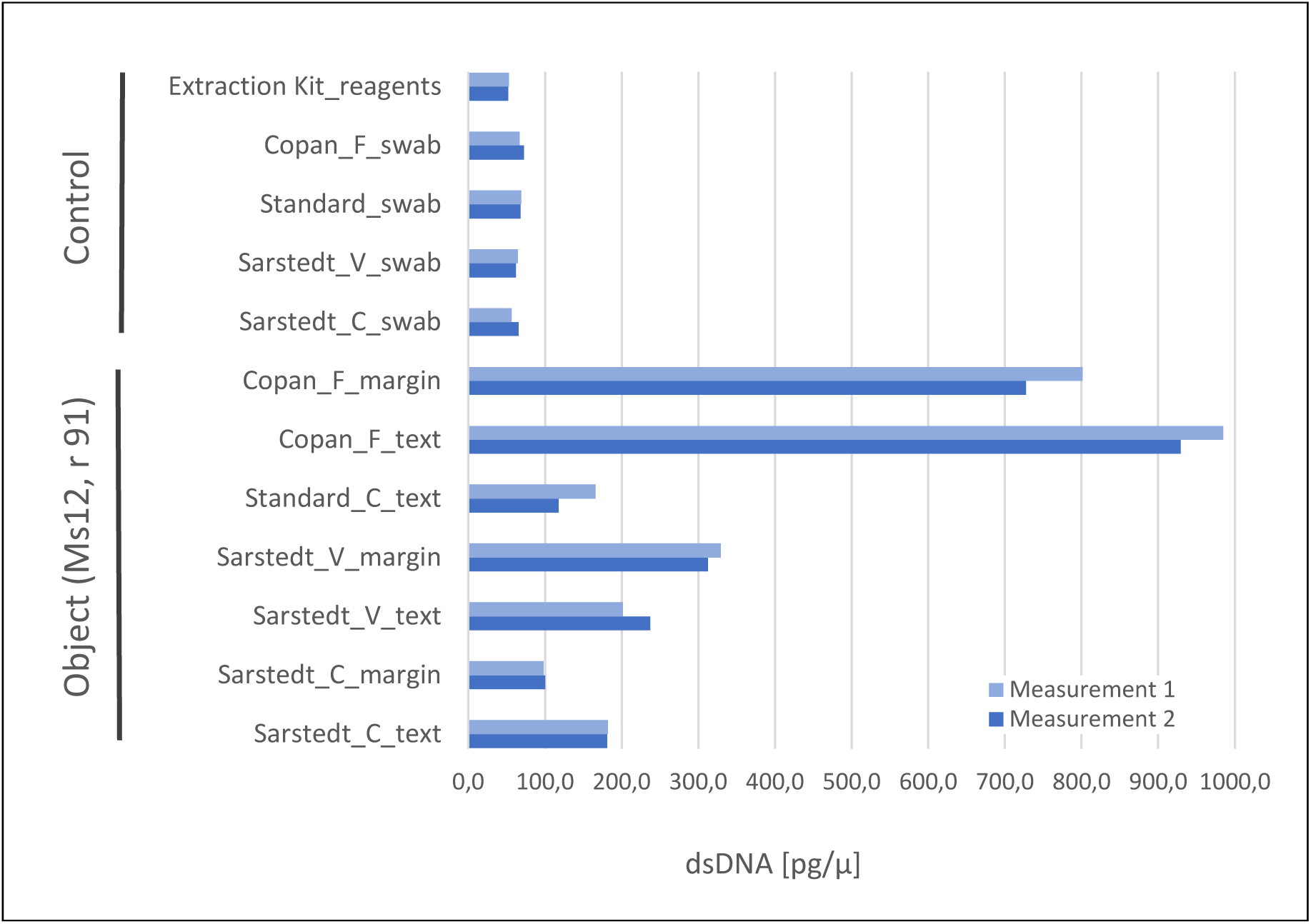
Comparison of the sampling performance of different types of swabs, with double stranded DNA (dsDNA) concentration as endpoint. One folium of a parchment codex (Ms12, 91r) was sampled in parallel with a set of different swabs, distinguishing between text and marginal areas with signs of use (each 100 cm^2^ surface), and subjected to nucleic acid extraction (QIAmp® DNA Micro Kit, Qiagen) and quantification (Quant-iT™ PicoGreen® dsDNA Kit, Invitrogen). For comparison purposes, all samples were taken from the same folium to avoid fluctuations bound to a potential different usage of the codex’s folia. Given the folium size constraints, one sample per experimental situation or control was taken, for which two technical replicates were generated for nucleic acids quantification. The combination of cotton swab and margin area was not produced due to the size limitation of the object. The controls consisted of swabs of each type without sample, which were extracted as the ones with sample, and also of extraction reactions without swab or sample, as control for extraction kit reagents. Abbreviations: material of the swab head (C) cotton, (V) viscose, (F) flocked nylon.

Considering all quantitative variables and hands-on experience, the nylon-flocked swabs were selected for downstream optimization tests and implementation into our standardized research protocols.

### 4.2 Optimizing the sampling procedure *on site*

For the sampling process itself, we devised a strategy based on the main procedures designed for microbial examination of flight and hardware in clean rooms by the European Cooperation for Space Standardization (ECSS, 2008), which we adapted and expanded for its application to cultural heritage objects. Basically, the original method starts by delineating a standardized sampling area from which a surface sample is taken with a sterile swab, previously soaked in a suitable buffer, under aseptic conditions. The swab head is carefully rolled over the surface of the object while moving forward on one direction till covering the whole sampling area. The process is repeated over the same area changing the swabbing motion by 90 and 135 degrees. Subsequently, the swab is stored or directly used to prepare a suspension for inoculation on solid cultivation medium (typically, a set with both, low and high nutrient content cultivation media), followed by incubation and colony counting after defined periods of time. Other pre-treatments (such has thermal or chemical shock) or selective cultivation media can be used to retrieve defined microbial taxa.

We recommend the application of the general steps of this standard methodology with the following modifications:

- Use a dry swab for sampling, since a moist one might not be suitable for certain cultural heritage objects. Also, the moistening process itself might increase the chances of contamination.
- Start the sampling by taking environmental control samples (for example, of the air surrounding the object) to avoid the dispersion of material and dust that may arise when sampling the object.
- Define the sampling areas in order to capture the heterogeneity of the object. Consider the dimensions of the object and the number of samples to be taken (for example, for our parchment codex we defined individual 100 cm^2^ areas).
- Prioritize taking biological/material replicates versus technical ones.
- Employ multiple cultivation media and conditions (e.g., atmosphere, solid/liquid, nutrient composition, pH, salinity), incubation times and selective pre-treatments, in order to capture a broader microbial diversity present on the object.

The final experimental setup will depend on the object under study. For example, for written heritage objects we recommend an initial set of cultivation media targeting non-fastidious and fastidious bacteria (such as R2A and tripticase soy yeast for the first, and Columbia blood agar for the latter group) and fungi that grow under high and low water availability conditions (for instance, malt extract agar and dichloran 18 % glycerol agar). The interactive database MediaDive (https://mediadive.dsmz.de/; Koblitz et al. 2022) offers a full catalogue of cultivation recipes and target microorganisms. With the described initial set of cultivation conditions, we were able to obtain a variety of cultivable microbiota present on the tested parchment codex. Stress resistant members of the *Bacillaceae*, including endospore-forming microorganisms, dominated among the isolates (Figure 5). Depending on the findings and additional lines of evidence that may emerge during the research, the set of cultivation conditions can be expanded to specifically target other microorganisms.

**Figure 5:**
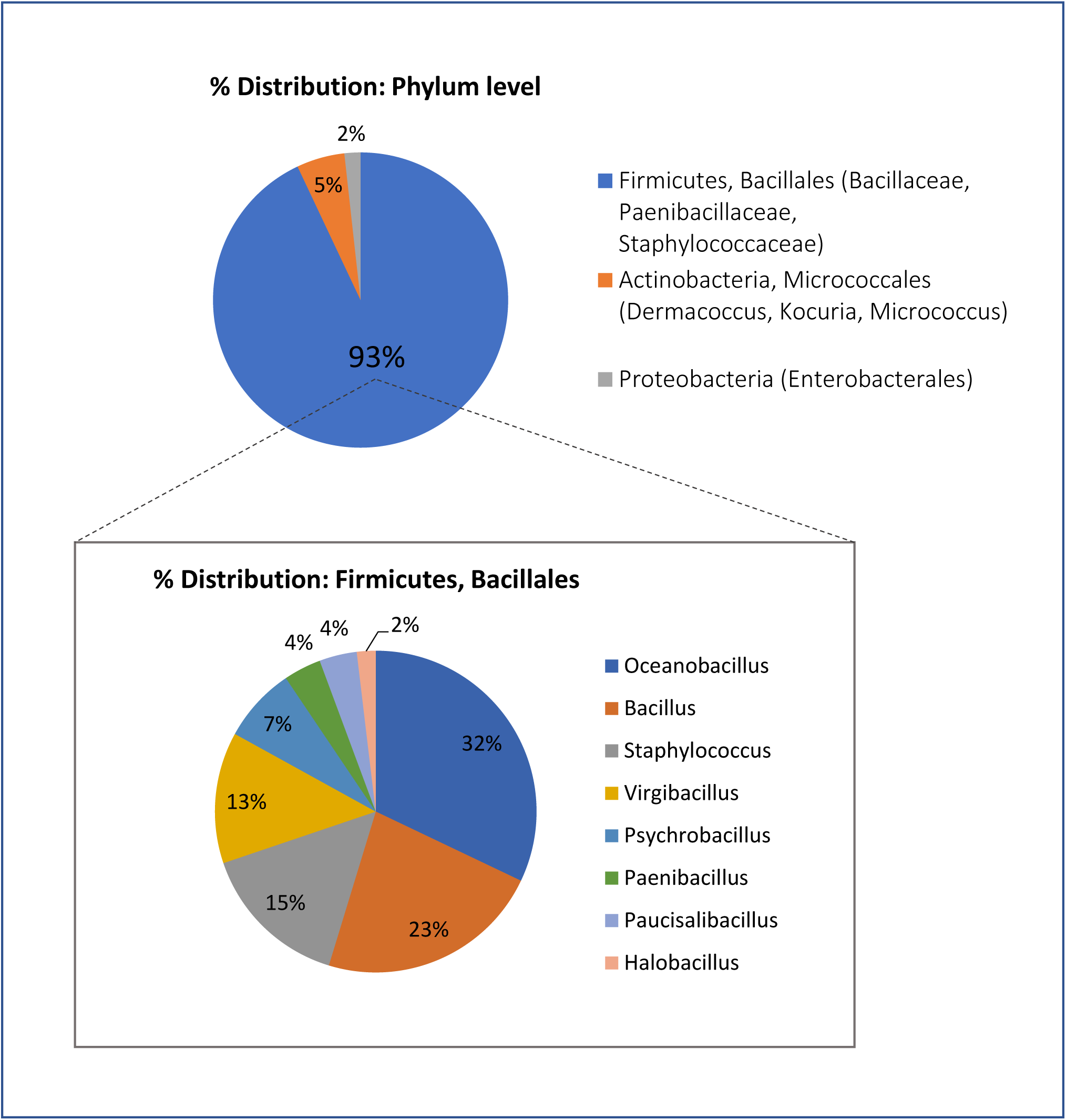
Distribution of microbial isolates retrieved from a parchment codex (Ms 12) used for the pilot study, phylum and genus level (16s rRNA gene-based classification, standard Sanger method).

To avoid contamination by the sampling process itself, the object should be moved, when possible, to a clean bench and work with appropriate aseptic technique and protective personal equipment, to avoid contamination by the operator. If this is not possible (for example when working with monuments or archive material that cannot be displaced or removed from its location), or when the institution hosting the object does not have laboratory facilities, the samples can be taken *on site* using appropriate protective personal equipment and pre-sterilized materials. The samples are immediately stored in appropriate containers to avoid further contamination (for example inside the dedicated transportation tube for swabs) and kept refrigerated at 4 °C (or at least, at a temperature similar to that at the location). Microbial cultivation steps should start as soon as possible, to avoid loss of viability of the microbial cells collected. If the samples are intended for cultivation independent analyses, a parallel set can be collected by using the same sampling procedure *on site* and kept frozen (at −20°C for DNA and −80°C for RNA analyses) till further processing.

### 4.3 Nucleic acids extraction and purification

Having sufficient starting material of good quality is essential for the sequencing process, therefore the extraction of nucleic acids is a critical step. Particularly, the surface samples of heritage objects may feature a low microbial biomass (and consequently, low content of nucleic acids of microbial origin) immersed a mixture of nucleic acids arising from different sources (Warinner et al., 2017). To tackle the optimization process, we carried out a literature-based approach combined with own wet lab assessments of the type of microorganisms that can be present in cultural heritage objects (Flocco and Overmann, 2021). Based on this combined approach, we preselected two commercial nucleic acids extraction kits for pilot testing. Although several in-home extraction methods exist (typically using a chloroform extraction), we focused on commercial nucleic acids extraction kits aiming to standardize the procedures, increase reproducibility, reduce the operational time and complexity, and cut the generation of toxic waste, which can be more difficult to handle at institutions without the required laboratory infrastructure. Custom made variations to the instructions of the manufacturers of commercial kits were only implemented when providing substantial advantages over standard procedures.

In addition to evaluating the main endpoint parameters, such as yield and integrity of the extracted nucleic acids, we focused our search on methods targeting samples with the above described low biomass and complexity characteristics. Such kits were developed for other types of samples (for example, low biomass medical samples such as lavages and tumours) but are potentially adaptable to cultural heritage objects. Also, we focused on kits designed to tackle samples rich in hard to lyse microbial cells, such as Gram positive and endospore-forming microorganisms that may constitute an important part of the cultural heritage microbiota, as previously mentioned.

Two commercially available nucleic acids extraction kits were identified as most promising. The QIAmp® DNA Micro Kit (Qiagen; protocol for isolation of genomic DNA from tissues), based on an enzymatic microbial lysis step and the DNeasy PowerBiofilm kit (Qiagen), which encompasses both enzymatic and mechanic lysis steps. All samples from the parchment manuscript used for this test were comparable and standardized by the surface area sampled (∼100 cm^2^) taken in duplicate. For both extraction methods, we also compared the operational performance and nucleic acids yield by directly incorporating the head of the sampling swab into the extraction workflow versus a previous release of the attached microbial cells and other materials into an extraction buffer and further extraction. In addition, we also tested the nucleic acids yield after one elution step and after sequentially extracting a second fraction, in order to maximize the amount of material collected. The experimental set up is shown in Figure 6. The extracted DNA was assessed with the Quant-iT™ Picogreen® dsDNA assay for an initial quantification and selected representative samples additionally tested with the ultrasensitive Femto Pulse system (Agilent), for yield, sizing and integrity. The QIAmp® DNA Micro Kit (Qiagen) yield was superior to that of the DNeasy PowerBiofilm kit (Qiagen) (∼2,4 ng/ul and 0,4 ng/ul, respectively, for a comparable surface sample area). The integrity of the nucleic acids extract was comparably well maintained, with both methods producing a high molecular weight peak. However, the DNeasy PowerBiofilm (Qiagen) which includes a mechanical lysis step, produced some low molecular weight fragments, as demonstrated by the sizing and quantification assay carried out with the ultrasensitive Femto Pulse system. (Figure 7).

**Figure 6:**
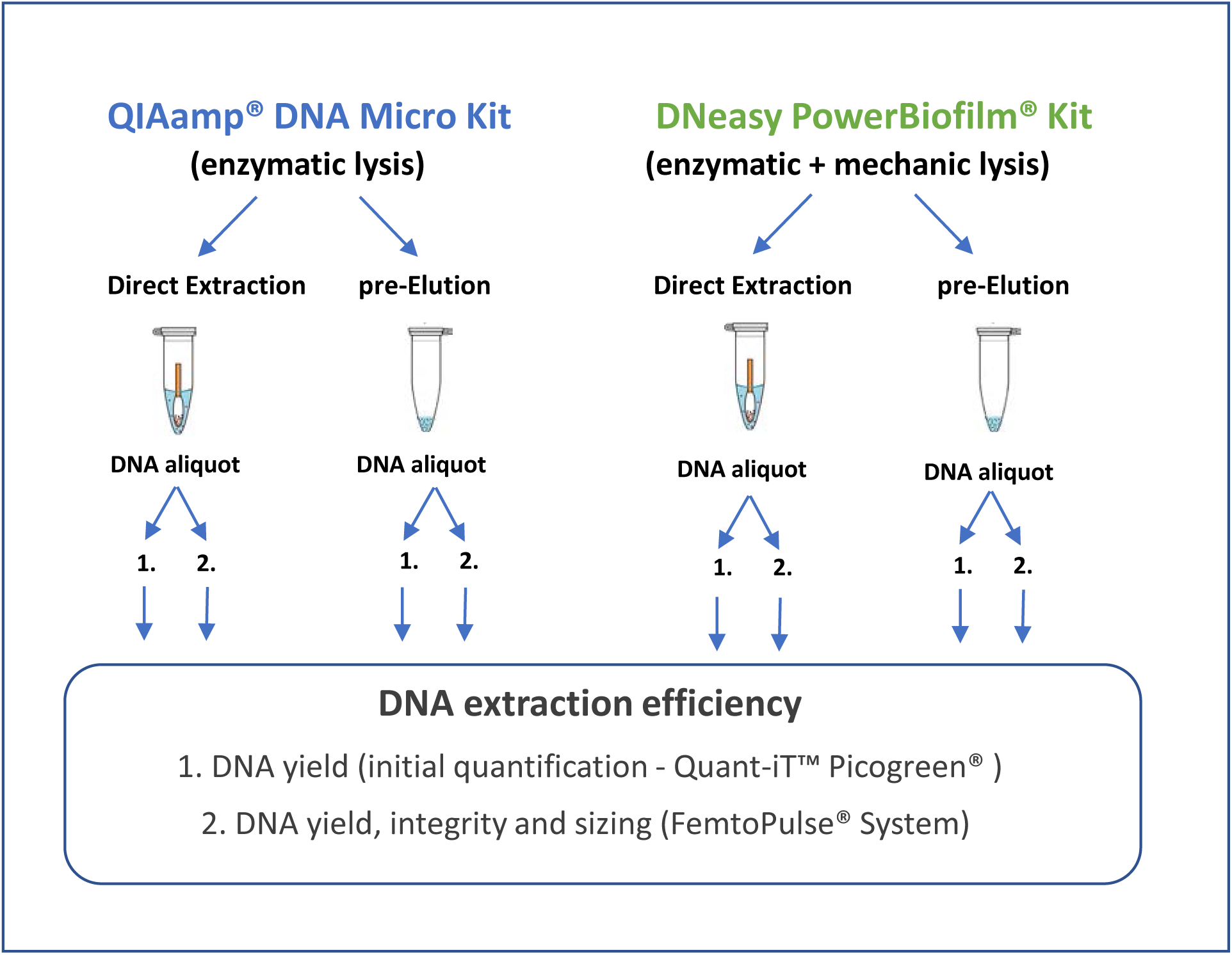
Schematic representations of the experimental design for the optimization of the methodology for nucleic acids extraction.

**Figure 7:**
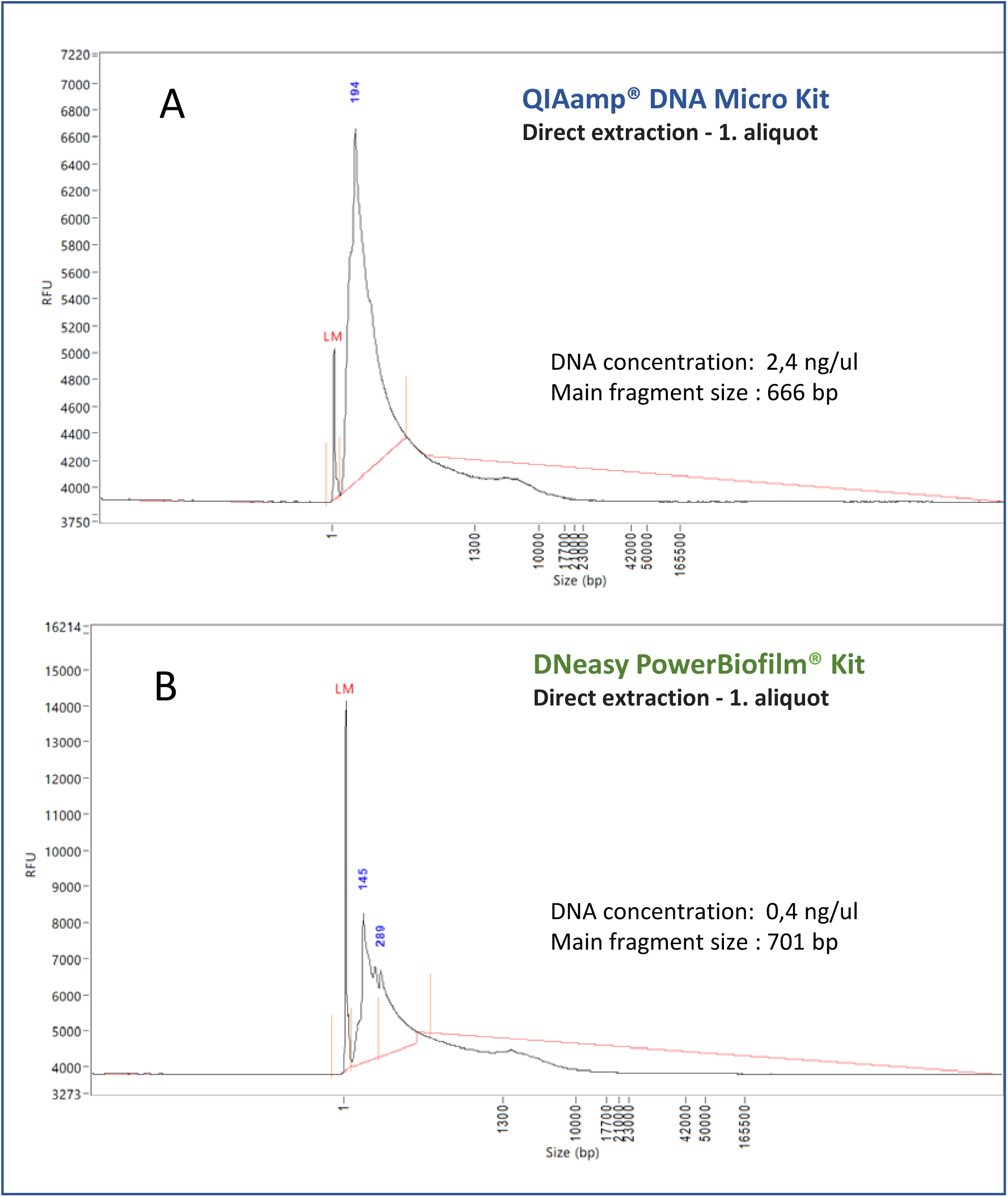
Quantification, integrity and sizing and of the nucleic acid extracts obtained through the methodology optimization process. Selected representative samples were analysed with the ultrasensitive Femto Pulse System (Agilent). (A) DNA extract profile obtained with the QIAmp® DNA Micro Kit (Qiagen), by direct extraction of the swab head and collection of one aliquot of nucleic acids extract (B) DNA extract profile obtained DNeasy PowerBiofilm kit (Qiagen) by direct extraction of the swab head and collection of one aliquot of nucleic acids extract.

Based on this pilot test we recommend:

- The QIAmp® DNA Micro Kit (Qiagen), selecting the protocol for tissue (with minor custom modifications, as indicated in the step-by step protocol included in this work).
- Start the extraction process by introducing the swab tip cut out directly into the extraction tube of the kit.
- Collect only one aliquot of DNA extract during the final clean up and elution step.

### 4.4 PCR amplification and sequencing library construction

A variety of possible combinations for the generation of an amplicon library of a phylogenetic marker are available, including the selection of the fingerprinting region(s) of the marker gene of choice (typically the hypervariable regions of 16S rRNA gene for prokaryotes and the internally transcribed spacer regions, ITS, for fungi; Schoch et al., 2012), several PCR amplification strategies, sequencing library preparation and loading protocols, sequencing platforms and run parameters, as well as the different steps of the bioinformatic pipeline. Here we focus on the bacterial community analyses using PCR amplification of the 16S rRNA gene and high throughput sequencing of amplicons. Due to the modular nature of the sequencing technology the individual workflow steps can be adapted to other taxa (see the step-by step protocol).

The selection of the target hypervariable region of the 16S rRNA gene is a key step, largely determined by the type of microbial taxa likely to be present in the objects to analyse and the aims of the study. Irrespective of the choice, it is important to bear in mind that an unavoidable degree of bias will exist, since amplification primers will have differential sensitivity toward microbial taxonomic groups (Abellan-Schneyder et al., 2021). Therefore, for comparison purposes, it is important to apply the same methodology across different studies. A literature survey can provide an overview of the type of microorganisms predominantly colonizing the object of study; in addition, inferences on microbial taxa occurrence can be made when considering the materiality and history of the object, which will provide the habitat for the microorganisms. If no previous information on the type of microbial entities colonizing the object of interest is available, or deductible, the target hypervariable region should provide as broad taxonomic information as possible, in order to capture most of the diversity present in the sample.

Once the target fingerprinting region is chosen, considerations about the sequencing approach itself should be made *a priori*, for example, assessing the desired sequencing depth and the generation of a single-end or paired-end amplicons. We recommend the latter approach since the target region is sequenced in both directions, forward and reverse. This allows the generation of a consensus sequence, which reduces the amount of ambiguous positions and the generation of artificial sequence diversity (produced by sequencing errors and not by true diversity). In addition, the generated amplicons should have a length (preferably around 300 base pairs) that allows a good taxonomic resolution, typically to the genus level or even down to species level, in some cases (Kullen et al., 2000)

Based on our synthesis of the literature on systematic benchmarking studies (Pollock et al., 2018; Abellan-Schneyder et al., 2021 and references therein) and own experimentation, we suggest to target the V3-V4 region, since this combination provides better taxonomic resolution, compared to other hypervariable regions of the 16S rRNA gene (Thijs et al., 2017; Abellan-Schneyder et al., 2021). Other hypervariable variable regions could be of interest, which might provide deeper taxonomic insight for specific taxonomic groups (Kullen at al., 2000).

Among the different primer combinations for targeting the V3-V4 region found in the literature, we highlight the options outlined below (targeting primarily bacteria), which can be adapted to better suit the research aims:

- The primer pair 347F - 803R (5’GGAGGCAGCAGTRRGGAAT 3’ - 5’CTACCRGGGTATCTAATCC3’) described by Nossa et al. (2010) has been developed for targeting bacteria with a focus on the human skin microbiome, which is an important feature when investigating heritage objects exposed to touch. This primer pair, originally implemented on a Roche 454 pyrosequencing platform (Nossa et al., 2010) has been later tested and validated for the Illumina MiSeq® platform (Castelino et al. 2017), which is a suitable sequencing technology for low biomass samples.
- Other useful primers for cultural heritage research, widely used and tested in large scale human and environmental microbiome studies, are the forward primer 341F (5’-CCTACGGGNGGCWGCAG-3’; Klindworth et al., 2013) and the reverse primer 806R (5’-GGACTACHVGGGTWTCTAAT-3’; Caporaso et al., 2011). These are described as universal (prokaryote) primers since they can detect both, Archaea and Bacteria.
- Alternatively, a custom modification of the forward primer 341F (5’-CCTACGGG<u>W</u>GGCWGCAG-3’; the underscore indicates the modification) has been shown to improve the detection of microbial taxa of soil and plant origin (Vieira et al., 2020), which can be of relevance for heritage objects exposed to the outdoor environment.

The selected 16S rRNA gene amplicon primers targeting the desired hypervariable region can be considered a modular and exchangeable component of a larger sequencing library primer construct, as they are concatenated with standard Illumina adapter sequences, binding sites for sequencing primers plus barcoding and index sequences for multiplexing. For generating a sequencing construct for the Illumina MiSeq® platform, we suggest a 2-step PCR amplification. The first PCR contains the 16S amplification primers for the selected hypervariable region (in this case the V4-V4 region) plus the barcode identifying each sample (attached to the forward primer) and the Illumina sequencing adapters (attached to the reverse primer). The second PCR provides the library index and Illumina adaptors. A full description is provided under the corresponding 16S Metagenomic Sequencing Library Preparation user’s manual (Illumina, 2013). This sequencing construct for the V3-V4 region produces an amplicon of ∼300 bp (this measure includes the target 16S amplicon plus the 16S sequencing primers and the barcode sequence; the latter two sections are trimmed later during the bioinformatic processing). Since it is sequenced in forward and reverse directions (paired-end approach) it provides a good overlap of the complementary sequences. Due to the modular approach employed, the same construct and sequencing strategy can be adapted to target other hypervariable regions by changing the 16S rRNA gene amplification primers, as discussed in previous paragraphs.

### 4.5 Bioinformatic Processing

After completion of the sequencing run, a series of sequencing data cleaning and trimming steps should be carried out before proceeding with microbial community analyses *per se*. The first step is to perform a quality check of the data, for example with the freely available FastQC package (Andrews, 2010) and discard those sequences that do not pass the desired quality parameters. If the sequences for a research project are obtained through more than one sequencing run, potential batch effects should be discarded before merging datasets. As Kennedy et al. (2023) highlight, it is recommended to use statistical approaches to determine the occurrence of batch effects and repeat some samples and negative controls across the different batches to help detect this effect.

The core microbiome bioinformatic analyses are typically carried out within bioinformatic platforms, such as Qiime2 (Boylen et al., 2019), which enable the execution of all steps under one roof. Following the initial cleaning of the output of the sequencing platform, the sequences are demultiplexed according to the barcode sequences assigned to each sample during the generation of the sequencing library; the accessory sequences of primers and adapters are trimmed. This is followed by a clustering step, in which the sequences are taxonomically classified and further grouped into operating taxonomic units (OTUs) or Amplicon Sequence Variants (ASVs). This is achieved using a classification pipeline and a taxonomic training set generated with a well curated and updated taxonomic database. Based on literature consensus on benchmarking studies (see reviews by Pollock at al., 2018; Abellan-Schneyder et al., 2021) and the experience collected in our team, we recommend implementing ASVs as clustering method and a well curated 16S rRNA database, such as SILVA (arb-silva.de; Quast et al., 2013) or the Ribosomal Database Project (RDP; rdp.cme.msu.edu), which maintain quality checked ribosomal RNA sequence data.

The ASVs count table can be further polished according to a set of bioinformatic parameters, excluding for example, samples which fall below a read count threshold or ASVs that have a percentual abundance below a given cut off value. Also, chimeras (artefacts formed during the sequencing procedure), sequences matching non-microbial references, such as human mitochondrial, or chloroplasts are further removed (although such type of sequences could be of interest, for example, for forensic studies or when working with heritage objects made with plant material). Processes and reagents contaminants can be bioinformatically removed. There are different approaches to executing this step, such as removing ASVs present in controls and sample datasets, but the procedure may discard true members of the community which are coincidentally also present as contaminants in reagents and procedures. Therefore, it is recommended to use statistical approaches to achieve this task. Kennedy et al. (2023) provide examples of this important post-sequencing step and suggested software packages.

Downstream statistical analyses are typically performed using R (R Core Team, 2020; www.r-project.org), a language and environment for statistical computing and visualization, and a menu of associated packages (such as *phyloseq* and *vegan*, and *ggplot* for graphs). These analyses provide microbial diversity metrics (such as alpha and beta diversity) and the possibility of linking the microbial community taxonomic and compositional information to the metadata associated to the sample under investigation (for example, source, material type, environmental conditions, age, location) with the help of multivariate analyses.

## 5 Conclusions and Final Recommendations

Over the last decades, the exploration of the interaction of microorganisms with cultural heritage objects has evolved, from the traditional perception that considers them agents of deterioration and disease, to a holistic approach which contemplates their potential historical, biographical and biotechnological value. The availability of high throughput sequencing platforms and improved microbial cultivation approaches has boosted such evolution, expanding cultural heritage research perspectives and opening new lines of inquiry of an interdisciplinary nature.

The inherent characteristics of cultural heritage objects (such as uniqueness, fragility, high value, restricted access) limit their availability for microbiological and biomolecular analyses, a situation that is exacerbated by the low microbial biomass present. This scenario requires the development of dedicated analytical and bioinformatical workflows and highlights the importance of a careful evaluation of upstream steps, such as the experimental design, the inclusion of critical controls and the sampling process.

The modular workflow proposed in this work was designed and optimized for its application to cultural heritage objects. Still, given the complex characteristics of these objects, several limitations should be considered:

- In many cases, the target heritage object is a unicate, which precludes the possibility of taking replicate samples and impacts the experimental design and downstream statistical analyses.
- All nucleic acids extraction methods have limitations, since they exhibit different degrees of efficiency largely associated to the resistance to lysis of the microbial cells and the presence of interferences (for example biofilms, sealing materials, substances binding to or degrading the nucleic acids, among other).
- The primers used for generating amplicons targeting different variable regions of 16S rRNA gene may introduce bias, since they have different sensitivity towards diverse taxonomic groups.
- Cultivation and cultivation-independent approaches provide different pictures of the microbial communities, with different degrees of taxonomic information overlap.
- Contemporary sources of microorganisms and nucleic acid may override older sources, a fact of particular importance when focusing on forensic and provenance research.
- The presence of microbial nucleic acids does not indicate the presence of viable microbial cells, an important consideration for the diagnosis of pathogens and biotic deterioration agents.

Where possible, mitigation measures to overcome these limitations should be applied. It may imply adaptation of wet lab protocols, *a priori* clean-up of contaminants and impurities and/or *a posteriori* optimization steps, such as *in silico* removal of spurious microbial signals and the optimization of the bioinformatic pipeline parameters. In spite of all precautions, some obstacles and deficits of the microbiome analysis cannot be sorted out. Accordingly, the inferences and conclusions derived from the experimental data should be crafted considering those limitations.

A prevalent challenge to the correct assessment of low biomass samples is the differentiation of legitimate data from contamination. As suggested for other microbiomes studies involving low microbial biomass samples (Kennedy et al. 2023), such critical assessment demands an interdisciplinary approach, which goes beyond the experimental work, to provide additional lines of evidence. In the case of cultural heritage studies, it may imply that hypothetical occurrence of certain microbial taxa is assessed considering as well environmental, historical, biographical and material aspects bound to the heritage object under study. Such holistic approach challenges the research hypothesis from different angles and reduces the chances of deriving erroneous conclusions or being impacted by the pervasive confirmation bias (Lange et al., 2021).

We intend this review to contribute towards expanding the perspectives of an interdisciplinary approach to cultural heritage research and to help identifying beforehand the main challenges it may pose. Aiming to bring theory into practice, we propose a step by step protocol in the following section, designed in a modular fashion to facilitate its adaptation to different cultural heritage scenarios and research needs.

## 6 Step by Step Protocol

### Experimental workflow for the study of the microbiome of heritage objects through 16S rRNA gene amplicon sequencing

#### Before starting

Timing: 2-4 days

Prepare in advance an accurate sampling plan. If photos or interactive digitized versions of the object under study are available, use those to aid the experimental design and delineation of the sampling scheme.

1. Calculate all necessary sampling materials and equipment (see Table 1)
2. Create sampling worksheets and pre-label materials.
3. Prepare all materials and mobile equipment for transportation (if sampling *on site*), observing requested storage temperatures and safety regulations for reagents and biological agents.

### 6.1 Materials and Equipment

**Table 1.**
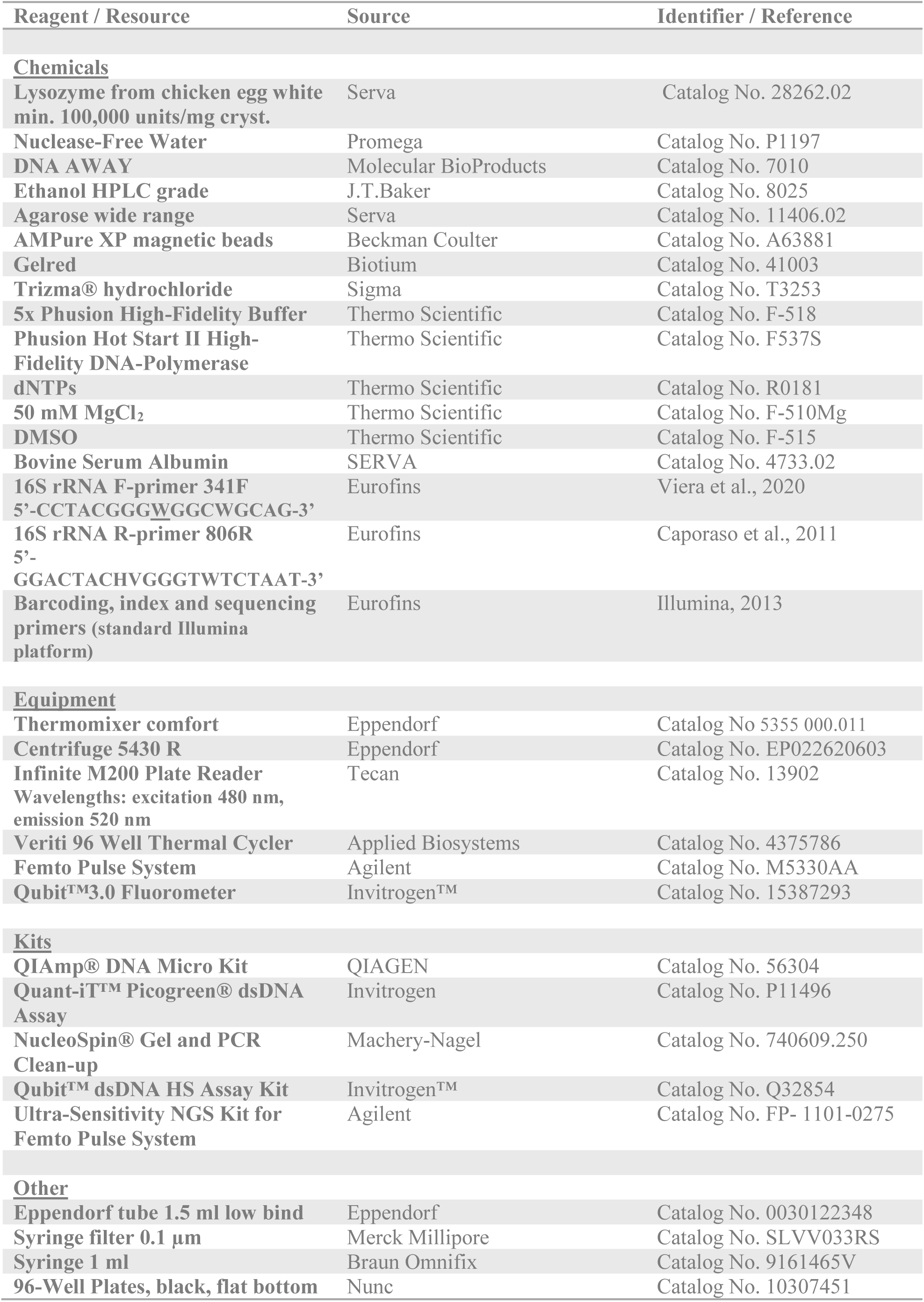
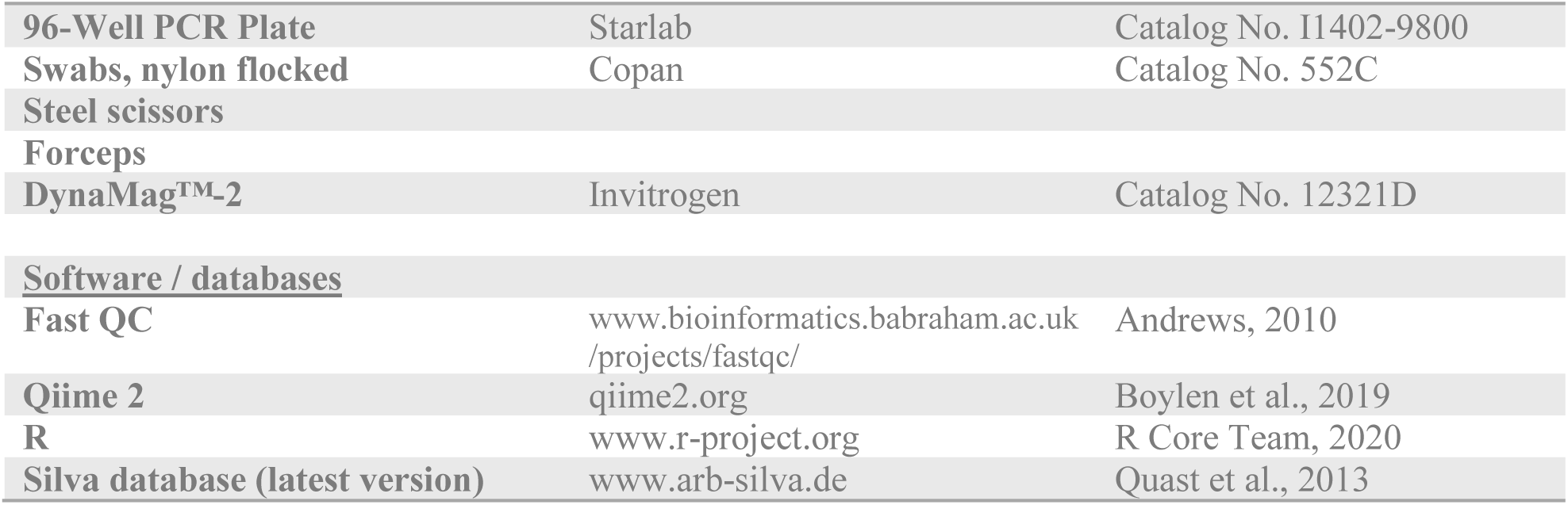
Key Resources

### 6.2 Step by Step Methodology

#### 6.2.1 Sampling and sample storage

Timing: 1h per 10 samples (sampling each 100 cm^2^ area).

**Note**: Transport the object of study to a clean bench. If not possible, minimize the exposure of the swabs to the environment. In all cases, use appropriate personal protection equipment and avoid shedding own skin, saliva drops or aerosols over the object; change gloves frequently.

1. Define the areas to be sampled. Place a ruler above of the object (without touching it) or use reference points on the object itself, to delineate the areas.
2. Take environmental control samples: expose a flocked swab to the environment surrounding the object under study (or to the clean bench area, if working under these conditions). Slowly rotate the swab to allow even distribution for a period of time similar to the one used for taking one sample of the object (next step)
3. Take samples from the object; carefully roll the swab head over the objects’ surface while moving forward on one direction till covering the whole sampling area. Repeated the process over the same area changing the swabbing motion by 90 and 135 degrees. Place the swab into its transportation tube and proceed with the next sample.
4. Store samples prior to DNA extraction at −-20 °C for DNA analyses. (or at −80 °C for RNA analyses). Freeze as well a set of not used swabs, which will serve as negative controls.
5. For cultivation experiments, initiate cultivation on site, if possible or store samples at 4 °C. Avoid extending the storage for more than 7 days, in order to preserver viability. We recommend direct plating on solid cultivation medium, but in some cases other pre-treatments might be necessary. The set of cultivation medium will depend on the goals of the study.

➔ Safe stop point

➔ DNA extraction should be performed in parallel for all samples including negative controls to minimize batch effects. If too many samples exist for a single batch, randomize the samples across different batches. Include a set of negative controls for reagents and procedures for each batch. When available also include some replicate object samples across batches, to aid the statistical analyses. Keep record of the kits lot number used for each batch of samples to track down eventual contaminations.

#### 6.2.2 DNA extraction from swabs

Timing: 8h (divided into 2 days: 4h on each day)

Before starting with the DNA extraction process:

1. Weigh 20 mg lysozyme on weighing tray wiped with DNA away, transfer to a sterile 1.5 mL centrifuge tube and add 1 ml nuclease-free water: Sterile filter the solution using a 0.1µm filter in a safety cabinet.
2. Prepare 10 mM Tris-HCl, pH 8,0 with nuclease-free water. Sterile filter the solution and treat it with UV overnight.

**Note:** Unless otherwise noted, all steps should be performed at room temperature (∼20°C). Page numbers mentioned in the protocol below refer to the QIAamp® DNA Micro Handbook 12/2014.

1. Set up safety cabinet and decontaminate with UV light. After this, wipe the surfaces with DNA Away.
2. Thaw frozen swabs.
3. Cut sterile head of swab using sterile scissors safety cabinet.
  a. Wipe steel scissors and forceps with DNA away.
  b. Dip them in 70% ethanol and flame them, wait for until tools cooled down.
  c. Cut the head of the swab into sterile low bind 1.5ml reaction tube. Avoid touching the border or exterior of the reaction tube with the swab head.
4. Preheat thermal block at 37°C.
5. Isolate DNA from the samples using the QIAamp® DNA Micro Kit according to the instructions of the manufacturer (QIAamp® DNA Micro Handbook 12/2014 Isolation of genomic DNA from tissues) working under the safety cabinet.
  a. Prepare buffers according to the instructions of the manufacturer instructions (p. 14-15).
  b. Add carrier RNA to Buffer AL (p.15).
  c. Add 180 µl Buffer ATL to the reaction tube containing the swab.
  d. Add 20 µl (20 mg/ml) lysozyme and pulse vortex ∼10 sec.
  e. Incubate for 1h at 37°C in thermal block, ∼500 rpm shaking.
  f. Add 20 µl Proteinase K and pulse vortex ∼10 sec.
  g. Incubate overnight at 56°C in thermal block, ∼500 rpm shaking. ➔ Overnight break
  h. Remove and discard the swabs from the tubes by using forceps wiped with DNA away, soaked in ethanol 100% and flamed. **Note:** Carefully pressing the swab head against the wall of the reaction tube is crucial for maximum DNA yield.
  i. Proceed with DNA extraction as described on step 5 use carrier RNA (p. 26)
  j. Elute in 1.5 ml low bind reaction tube by applying 2 x 10 µl nuclease-free water to the column, incubate loaded column each time for 5 min at room temperature prior to centrifugation.

➔ Safe stop point. DNA can be stored at −20°C for further applications.

#### 6.2.3 DNA quantification

Timing: ∼ 4h for 60 samples including analysis.

DNA yield is quantified using the Quant-iT™ PicoGreen ® dsDNA Kit according to the instructions of the manufacturer for standard procedure (https://www.thermofisher.com/document-connect/document-connect.html?url=https%3A%2F%2Fassets.thermofisher.com%2FTFS-Assets%2FLSG%2Fmanuals%2Fmp07581.pdf&title=UXVhbnQtaVQgUGljb0dyZWVuIGRzRE5BIFJlYWdlbnQgYW5kIEtpdHM=) Volumina are reduced for high-throughput assay.

A total volume of 200 µl per assay is prepared in a flat bottom black 96-well plate.

**Note**: if working with very low yield DNA samples, it is recommended to use the Femto Pulse System.

1. Allow the Quant-iT™ PicoGreen® reagent to warm to room temperature before usage.
2. Make sure fluorescence microplate reader is available with standard fluorescein wavelengths (excitation ∼480 nm, emission ∼520 nm).
3. Prepare the sterile TE buffer (10 mM Tris-HCl, 1 mM EDTA, pH 7.5). Using the included 20x TE a 1x TE working solution is prepared by diluting the concentrated buffer 20-fold with nuclease-free water.
4. Prepare the PicoGreen® working solution by dissolving PicoGreen® reagent 1:200 in 1x TE. **Note:** Use a plastic tube, as the reagent may adsorb to glass. Protect the PicoGreen® working solution from light, it is susceptible to photo degradation. Use it immediately.
5. Dilute provided lambda DNA standard (100 µg/ml) 50-fold in 1x TE to make a final high-range stock solution of dsDNA (2 μg/mL).
6. Prepare a low-range stock solution of dsDNA (50 ng/ml) by diluting the high-range solution 40-fold.
7. Prepare high-range and low-range standard curves as shown in table 2 in 96-well plate, black, flat bottom.
8. Add 100 µl PicoGreen® working solution to all wells.
9. Pipette 1 µl of DNA extract in 96-well plate.
10. Dilute each DNA 100-fold by adding 99 µl 1x TE buffer.
11. Add 100 µl PicoGreen® working solution to the diluted extracts.
12. Incubate plate for 2-5min in the dark (cover it with aluminum foil).
13. Measure the fluorescence 480 nm (excitation) and 520 nm (emission) with fluorescence microplate reader.
14. Subtract the fluorescence value of the reagent blank from that of each sample.
15. Create a standard curve and determine the DNA concentration of all samples accordingly.

**Table 2.** Calculation of PicoGreen® solutions

| Low-range<br>DNA<br>Standards | final conc.<br>(pg/μl) | TE (1x)<br>[μl] | DNA Stock<br>(50 ng/ ml)<br>[μl] | PicoGreen®<br>Working<br>Solution<br>[μl] |
| --- | --- | --- | --- | --- |
|  | 25.0 | 0.0 | 100.0 | 100.0 |
|  | 2.5 | 90.0 | 10.0 | 100.0 |
|  | 0.25 | 99.0 | 1.0 | 100.0 |
|  | 0.025 | 99.9 | 0.1 | 100.0 |
| Blank | 0.0 | 100.0 | 0.0 | 100.0 |
|  | Total [μl] | 388.9 | 111.1 | 500.0 |
| High-range<br>DNA<br>Standards | final conc.<br>(pg/μl) | TE (1x)<br>[μl] | DNA Stock<br>(2 μg/ ml)<br>[μl] | PicoGreen®<br>Working<br>Solution<br>[μl] |
|  | 1.0 | 0.0 | 100.0 | 100.0 |
|  | 0.1 | 90.0 | 10.0 | 100.0 |
|  | 0.01 | 99.0 | 1.0 | 100.0 |
|  | 0.001 | 99.9 | 0.1 | 100.0 |
| Blank | 0.0 | 100.0 | 0.0 | 100.0 |
|  | Total [μl] | 388.9 | 111.1 | 500.0 |

**Note:** Minimize photo bleaching effects by keeping the time for preparation and fluorescence measurement constant for all samples

➔ Safe stop point. Freeze DNA at −20°C

#### 6.2.4 Library preparation by PCR amplification of 16S rRNA gene V3-V4 region

Timing: ∼ 4h for 20 samples

The V3-V4 region of the 16S rRNA gene is amplified using primers 341-F 2w and 803-R primers. The forward primer contains a 6-nt barcode and the reverse primer contains a 6-nt index. Both primers comprise sequences complementary to the Illumina specific adaptors to the 5′-ends. (for full details please see the library preparation guide under Illumina, 2013). Library preparation is based on a two-step PCR.

- **Note:** preparation may be performed in a UV-sterilized biosafety hood. Pay careful attention to pipetting precision. Other amplicon primer pairs can be coupled to the Illumina primers, according to the study aims.

1^st^ PCR:

1. Prepare master mix according to Table 3, add 18 µl to each well of the 96-well plate.
2. Add 0.5 µl of specific forward primer barcode to 96-well plate, each well gets a different one.
3. Transfer 3 µl of DNA into the corresponding well.
4. Mix total volume of 21.5 µl by pipetting up and down.
5. Seal the plate with foil plate covers and perform amplification in the thermocycler, use cycle conditions stated in table 5a.

**Table 3.**
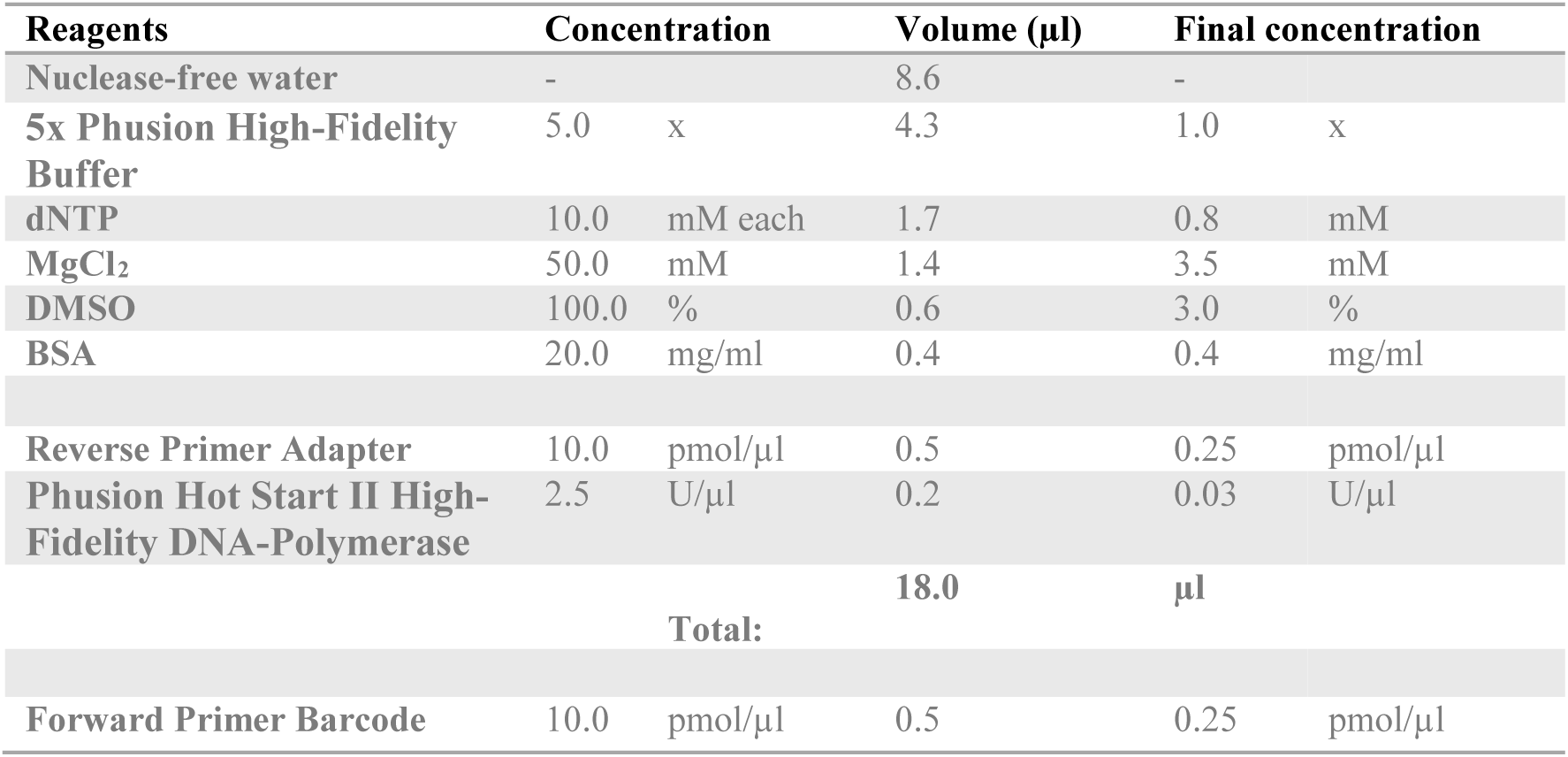
PCR master mix 1st PCR

2^nd^ PCR:

1. Prepare master mix according to table 4 for each reverse primer index you like to use, add 50 µl to each well of the 96-well plate.
2. Transfer 2 µl of PCR product of 1^st^ PCR to according wells. Each sample is prepared in triplicates.
3. Non-template controls are performed using same conditions.
4. Mix total volume of 52.0 µl by pipetting up and down.
5. Seal the plate with foil plate covers and perform amplification in the thermocycler, use cycle conditions stated in table 5b.

**Table 4.**
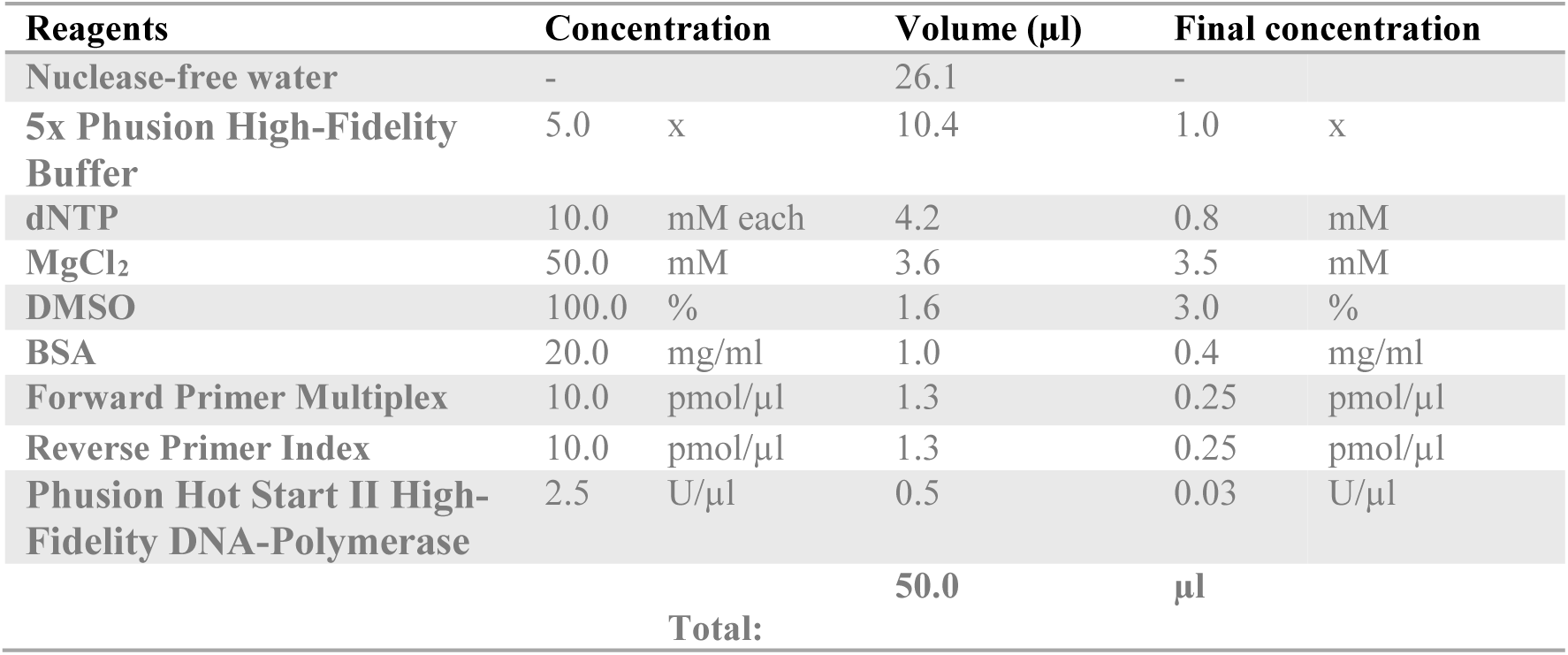
PCR master mix ^2nd^ PCR

➔ PCR amplicons were verified by gel electrophoresis stained with GelRed (6 µl Gelred/100ml 1% agarose).

**Table 5a.** PCR cycling conditions 1^st^ PCR

|  |  | T (°C)/ Time |
| --- | --- | --- |
| 1 | Denaturation | 98/ 30 sec |
| 2 | Denaturation | 98/ 10 sec |
| 3 | Annealing | 55/ 10 sec |
| 4 | Elongation | 72/ 45 sec |
|  | Cycle: Step 2-4 | 20x |
| 5 | Final elongation | 72/2min |
| 6 | End | 4/∞ |

**Table 5b.** PCR cycling conditions 2^nd^ PCR

|  |  | T (°C)/ Time |
| --- | --- | --- |
| 1 | Denaturation | 98/ 30 sec |
| 2 | Denaturation | 98/ 10 sec |
| 3 | Annealing | 55/ 10 sec |
| 4 | Elongation | 72/ 45 sec |
|  | Cycle: Step 2-4 | 15x |
| 5 | Final elongation | 72/ 2 min |
| 6 | End | 4/ ∞ |

#### 6.2.5 Purification of library products

Timing: ∼ 1. 5 h for 20 samples

Using Macherey-Nagel kit following the instructions of the manufacturer for clean-up of PCR products (see page 17/18 Protocols, PCR clean-up; https://www.mn-net.com/media/pdf/02/1a/74/Instruction-NucleoSpin-Gel-and-PCR-Clean-up.pdf).

**Note:** PCR amplicon triplicates were pooled for clean-up.

1. Mix 1 volume of sample with 2 volumes of binding buffer NTI: 150 µl PCR product + 300µl NTI
2. Place a NucleoSpin Gel and PCR Clean-up column into a collection tube (2ml) and load up to 700 µl sample.
3. Centrifuge for 30 s at 11,000 x g. Discard flow-through and place the column back into the collection tube.
4. Load remaining sample if necessary and repeat the centrifugation step.
5. Add 700 µl wash buffer NT3 to the NucleoSpin® Gel and PCR Clean-up column. Centrifuge for 30s at 11,000 x g. Discard flow-through and place the column back into the collection tube.
6. Repeat previous washing step.
7. Centrifuge for 1 min at 11,000 x g to remove wash buffer NT3 completely. Make sure the spin column does not come in contact with the flow-through while removing it from the centrifuge and the collection tube. **Note:** Residual ethanol from wash buffer NT3 might inhibit enzymatic reactions. Total removal of ethanol can be achieved by incubating the columns for 2-5 min at 70°C prior to elution.
8. Place the NucleoSpin® Gel and PCR Clean-up column into a new 1.5 ml microcentrifuge tube. Add 15 µl nuclease-free water for elution and incubate at room temperature for 1 min. Centrifuge for 1 min at 11,000 x g. Repeat previous elution step.

#### 6.2.6 Quantification of libraries using the Qubit™ dsDNA HS Assay Kit

Timing: ∼ 0.5 h for 20 samples

1. Prepare the Qubit™ working solution by diluting the Qubit™ dsDNA HS Reagent 1:200 in Qubit™ dsDNA HS Buffer. Use a clean plastic tube each time you prepare Qubit™working solution. **Note:** Do not mix the working solution in a glass container. The final volume in each tube must be 200 μL. Each standard tube requires 190 μL of Qubit™working solution, and each sample tube requires anywhere from 180–199 μL. Prepare sufficient Qubit™working solution to accommodate all standards and samples.
2. Add 190 μL of Qubit™ working solution to each of the tubes used for standards.
3. Add 10 μL of each Qubit™ standard to the appropriate tube, then mix by vortexing 2– 3 seconds. Avoid producing bubbles. ***Note:*** *Careful pipetting is critical to ensure that exactly 10 μL of each Qubit*™ *standard is added to 190 μL of Qubit*™ *working solution*.
4. Add Qubit™ working solution to individual assay tubes so that the final volume in each tube after adding sample is 200 μL. **Note:** Your sample can be anywhere from 1–20 μL. Add a corresponding volume of Qubit™working solution to each assay tube: anywhere from 180–199 μL.
5. Add each sample to the assay tubes containing the correct volume of Qubit™ working solution, then mix by vortexing 2–3 seconds. The final volume in each tube should be 200 μL.
6. Allow all tubes to incubate at room temperature for 2 minutes. Proceed to “Reading standards and samples”; follow the procedure appropriate for your instrument “Qubit™ 3.0 Fluorometer”

#### 6.2.7 Pooling equimolar libraries for sequencing

Timing: ∼ 0.5 h for 20 samples

1. Calculate molarity of each sample using the dsDNA concentration and fragment size (550 bp for the procedures described I this protocol). The described variables can be introduced directly on an online calculator (for example, www.promega.de/resources/tools/biomath), which is based on the following formula:

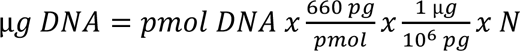

where
N = DNA fragment length, in bp (550 bp, in this case) 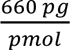 = average molecular weight of a nucleotide pair 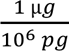 = units conversion factor
2. Dilute samples with resuspension buffer to get equimolar samples.
3. Take 5 µl of each diluted sample and mix properly in a microcentrifuge tube.

**Note:** In case of very low amount of library concentration it is recommended to concentrate the pool.

#### 6.2.8 Clean up and concentration of DNA library pool with magnetic beads

Timing: ∼ 0.5 h

**Aim:** elimination of small side-products and concentration of library pool

**Note:** All steps are performed at room temperature (∼ 22 ◦C)

1. Allow AMPure XP magnetic beads to warm to room temperature before usage.
2. Vortex beads for 30 seconds.
3. Add 60 µl beads to 60 µl pooled PCR products, mix by pipetting (carefully 10x).
4. Incubate for 5 min.
5. Put microcentrifuge tube on magnetic rack (DynaMag™-2) and wait for 2 min. The supernatant is supposed to get clear. **Note:** For step 4-7 keep microcentrifuge tube on magnetic rack.
6. Discard the supernatant.
7. Wash beads with 200 µl 80% ethanol, incubate for 30 s and discard supernatant.
8. Repeat previous step.
9. Remove residual ethanol by using thin 10 µl tips and dry for 2 min.
10. Take microcentrifuge tube of magnetic rack and add 30 µl resuspension buffer (10 mM Tris-HCl, pH 8.0) and mix by pipetting gently.
11. Incubate for 5min.
12. Put microcentrifuge tube on magnetic rack and wait 2 min. The supernatant is supposed to get clear.
13. Transfer supernatant containing cleaned-up library pool into new tube.

#### 6.2.9 Quantitation of library pool using the Qubit™ dsDNA HS Assay Kit

**Aim:** this step is preceding the quantitation and sizing of the library using the ultrasensitive Femto Pulse system. It allows the estimation of the sample concentration and the calculation of the dilution factor to be applied to the sample for its analysis with Ultra Sensitivity assay for Femto Pulse system.

➔ see previous section for Qubit™ Assay

#### 6.2.10 Quantitation and sizing of the library pool using Agilent Femto Pulse System

Timing: ∼ 1.5 h

Use the Ultra Sensitivity NGS kit for Femto Pulse System for quantitative and qualitative analysis of NGS libraries according to the instructions of the manufacturer (https:\\www.agilent.com\\cs\\library\\usermanuals\\public\\quick-guide-fp-1101-ultra-sensitivity-ngs-kit-SD-AT000143.pdf) Before starting, review the instructions for the preparation and storage of reagents and machine set up and maintenance (see pages 4-5 for sample preparation and general instructions for set up and running).

**Note:** The total input DNA in the sample must be within a range of 0.1 pg/ul to 5 pg (DNA fragment) or 5 pg/µl to 250 pg/µl (DNA smear). Estimate the sample concentration using the Qubit**™** assay, if necessary, dilute the sample with 0.25 TE buffer before performing the Femto Pulse assay.

1. Pipette 18 µl of diluted Diluent Marker (DM) solution (prepared according to manufacturer’s Instructions, as indicated above) into each well of a well-sample plate that will carry sample. Fill unused wells with 20µl of blank solution (kit reagent BF-P25)
2. DNA ladder (prepared according to instructions of the manufacturer, as indicated above) is run in parallel with the samples. Pipette 2 µl of ladder into the well number 12 of each row to be analyzed and mix well by swirling while aspiring and expulsing with the pipette tip.
3. Load 2 µl of sample to the well containing 18 µl of DM solution and mix well as indicated above.
4. After all loading steps, place a plate seal, vortex at 3000 rpm for 2 minutes and then centrifuge the plate to remove air bubbles; check the bottom of the wells to ensure this condition.
5. Place the plate on one of the sample trays of the Femto Pulse instrument and load the experimental method and follow the software operational procedure as described in the instructions of the manufacturer (page 7)
6. A satisfactory sample should show a peak for the product size expected for the chosen 16S rRNA gene amplicon library and an absence of secondary peaks of other sizes.

#### 6.2.11 Denaturation and dilution of the library for sequencing in MiSeq® System (Illumina®)

Timing: ∼ 0.5 h

The protocol A to denature and dilute libraries normalized standard library quantification is used. In addition, a protocol modification for sequencing low biomass and low abundance samples on a MiSeq**®** System (Illumina®) for a 2 nM library was used. The denature and dilution steps are compatible with the MiSeq**®** Reagent kit v2. The main steps are indicated below (please refer to the manufacturer’S handbook for other protocol options and additional recommendations; https://support.illumina.com/content/dam/illumina-support/documents/documentation/system_documentation/miseq/miseq-denature-dilute-libraries-guide-15039740-10.pdf)

1. Prepare a fresh dilution of NaOH 0.2 N by combining 800 µl of laboratory grade water and 200 µl of a stock solution of 1.0 N NaOH. Chill reagent HT1 (included in kit). **Note:** NaOH dilution should be prepared freshly and used within 12 hours. Preferably prepare a large volume (1 mL) to minimize pipetting errors.
2. Denature a 2.0 nM library: Combine 5 µl of the 0.5 nM library with 5 µl of 0.2 N NaOH in a microcentrifuge tube.
3. Vortex at 280 x g for 1 minute and incubate at room temperature for 5 minutes.
4. Add 990 µl of prechilled reagent HT1 to the microcentrifuge tube, to produce 1 mL of a 10 pM library.
5. Use the library at that concentration or dilute the 10 pM library to the desired concentration with prechilled HT1a reagent: for example, mix 360 µl of 10 pM library with 240 µl of HT1 reagent to produce a 6 pM library. Mix by inversion of the tube and pulse centrifuge.

➔ Safe stop point: seal the plate (avoiding well cross-contamination) and store at −25°C to −15°C.

#### 6.2.12 Denaturation and dilution of the Phi X control library for sequencing in MiSeq® System (Illumina®)

Timing: ∼ 0.5 h

1. Dilute the 10 nM PhiX library (included in kit) to 4 nM by combining 2 µl of PhiX library with 3 µl of Tris-Cl, pH 8.5 with 0.1% Tween 20.
2. Denature the 4 nM Phi X library: in a microcentrifuge tube combine 5 µl of the 4 nM library with 5 µl of 0.2 N NaOH (prepared within the last 12 hours).
3. Vortex at 280 x g for 1 minute and incubate at room temperature for 5 minutes.
4. Add 990 µl of prechilled reagent HT1 to the microcentrifuge tube, to produce 1 mL of a 20 pM library.

#### 6.2.13 Spike the test library with PhiX control library

The protocol of the manufacturer for spike in for low diversity libraries (recommended >5% spike in; page 11) was modified as follows:

1. Combine 540 µl of diluted and denaturated test library (10 pM) with 60 µl of diluted and denatured PhiX Control library (20 pM). This yields a PhiX concentration of 10%.
2. Mix by pulse vortex and spin down. Load onto the MiSeq**®** reagent cartridge (or set on ice until ready to load) and set up the sequencing run (600 cycles generating paired-end sequences with 300 base pairs per read) according the MiSeq® user guide (https://support.illumina.com/content/dam/illumina-support/documents/documentation/system_documentation/miseq/miseq-denature-dilute-libraries-guide-15039740-10.pdf)

#### 6.2.14 Bioinformatic Analysis

Here we provide a succinct guideline to the main steps of the bioinformatic pipeline and recommended software and literature. The final workflow will depend on the type of samples, chosen sequencing approach and study aims.

1. After completion of the sequencing run, check the quality of the raw sequences (usually provided in FASTQ format) to broadly detect common sequencing contaminants (for example, with FastQC package; Andrews, 2010).
2. Check for batch effects if more than one sequencing run was used. This can be done by comparing negative controls and a set of samples run in all sequencing batches. Dedicated statistical approaches can be implemented for this step (Kennedy et al. 2023). Discard SV not consistently detected along all batches before merging datasets.
3. Demultiplex the reads according to the barcodes and trim sequences belonging to primers and adapters. The core sequence data analyses can be carried out with the microbiome bioinformatic platforms Qiime2 (Boylen et al., 2019),
4. Generate an amplicon sequence variant (ASV) table using the most recent SILVA (www.arb-silva.de; Quast et al., 2013) taxonomic dataset trained for the 16S rRNA gene variable regions/s selected for the study.
5. Standardize the datasets by establishing cut offs, selected according to the different research needs (for example exclude samples with less 1000 reads, include SVs with more than 1% abundance in any sample and other parameters.
6. Prune: eliminate SV corresponding to human mitochondrial sequences and chloroplasts and eventually taxa present in controls. For the latter it is best to use a statistical package (see Kennedy et al., 2023) to avoid eliminating low abundance community members, which could be confounded with contaminants.
7. Downstream analyses are typically carried out with the R environment for statistical computing (R Core team, 2020) and includes diversity metrics and multivariate analyses.

### 6.3 Troubleshooting

**Challenge**: some samples carrying very low biomass, and/or low diversity may not yield enough amplicon product for sequencing or may generate a number of reads falling below quality filters and have to be discarded.

**Solution**: this challenge can be tacked at different steps of the library generation and sequencing protocol:

a. Generation of PCR products: produce multiple PCR runs for each sample and pool them, to increase the total amount of amplicon per sample.
b. Concentration of the library pool: modify the elution volume of the purification and concentration steps with AM Pure beads, to increase the concentration of the library product.
c. Modify the library denaturation and dilution steps for low biomass, low diversity samples, which typically works with a 2 nM library, as follows:
  1. Start working with a 0.5 nM library (instead of the 2.0 nM library described in the protocol)
  2. Denature the library library by combining 20 µl of the 0.5 nM library with 20 µl of 0.2 N NaOH.
  3. Vortex at 280 x g for 1 minute and incubate at RT for 5 minutes.
  4. Add 960 µl of prechilled of reagent HT1 to the microcentrifuge tube, to produce 1 mL of a 10 pM library.
  5. Spike 540 µL of the denatured and diluted 10 pM library with the denatured and diluted PhiX library 20 pM (prepared as previously described). This yields a denatured and diluted library spiked with PhiX control at 10 % ready to load on the MiSeq® Instrument.

**Challenge**: DNA extraction fails, produces degraded DNA.

**Solution**: the addition of β-mercaptoethanol, a reducing agent, to the lysis buffer of the nucleic acids extraction protocol (0.2-0.5 % vol/vol) may help prevent oxidative damage and nuclease activity (Healey et al., 2014).

## 8 Author Contributions

CF and JO conceived the study. JO provided funding. CF designed the workflow and performed the sampling, experiments and data analyses. AM, FB and AG performed experiments. CF wrote the manuscript. AM, FB contributed writing to the step-by-step protocol. All authors contributed to manuscript revision, read, and approved the submitted version.

## 10 Acknowledgments

The authors would like to thank Dr. Christoph Mackert and Prof. Ulrich Schneider (Manuscript Center and University Library Leipzig, Germany) for kindly providing access to the written cultural heritage objects used for optimizing the proposed protocol; Dr. Manfred Rohde for his help to create the SEM image; Dr. Selma Vieira for providing information on the 16S primers and Dr. Boyke Bunk, Dr. Cathrin Spröer and the team at sequencing facility at the Leibniz Institute DSMZ for their support on the sequencing procedures.

## 11 Data Availability Statement

The original research contributions are included directly in this article; further inquiries can be directed to the corresponding author.

